# Decoding Gene Networks Controlling Hypothalamic and Prethalamic Neuron Development

**DOI:** 10.1101/2025.01.10.632449

**Authors:** Dong Won Kim, Leighton H. Duncan, Jenny Xu, Minzi Chang, Sara Sejer Sørensen, Chantelle E. Terrillion, Patrick O. Kanold, Elsie Place, Seth Blackshaw

## Abstract

Neuronal subtypes derived from the embryonic hypothalamus and prethalamus regulate many essential physiological processes, yet the gene regulatory networks controlling their development remain poorly understood. Using single-cell RNA- and ATAC-sequencing, we analyzed mouse hypothalamic and prethalamic development from embryonic day 11 to postnatal day 8, profiling 660,000 cells in total. This identified key transcriptional and chromatin dynamics driving regionalization, neurogenesis, and differentiation. This identified multiple distinct neural progenitor populations, as well as gene regulatory networks that control their spatial and temporal identities, and their terminal differentiation into major neuronal subtypes. Integrating these results with large-scale genome-wide association study data, we identified a central role for transcription factors controlling supramammillary hypothalamic development in a broad range of metabolic and cognitive traits. Recurring cross-repressive regulatory relationships were observed between transcription factors that induced prethalamic and tuberal hypothalamic identity on the one hand and mammillary and supramammillary hypothalamic identity on the other. In postnatal animals, *Dlx1/2* was found to severely disrupt GABAergic neuron specification in both the hypothalamus and prethalamus, resulting in a loss of inhibition of thalamic neurons, hypersensitivity to cold, and behavioral hyperactivity. By identifying core gene regulatory networks controlling the specification and differentiation of major hypothalamic and prethalamic neuronal cell types, this study provides a roadmap for future efforts aimed at preventing and treating a broad range of homeostatic and cognitive disorders.

## Introduction

In the embryonic forebrain, the anterior diencephalon consists of the hypothalamus and prethalamus, both essential for various neurophysiological functions^1–4^. The hypothalamus, which contains hundreds of molecularly distinct neuronal subtypes^5^, regulates a broad range of homeostatic physiological processes and innate behaviors, including circadian rhythms, the sleep/wake cycle, food intake, reproduction, and stress response^6,7^. In contrast, the prethalamus gives rise to the zona incerta (ZI) and thalamic reticular nucleus (TRN), which almost entirely consist of GABAergic neurons^8,9^. GABAergic neurons in the ZI and TRN are key regulators of sensory gating^10–12^, inhibiting targets in the cerebral cortex and dorsal thalamus^1–4^. The ZI has recently been found to regulate many homeostatic behaviors historically attributed to the hypothalamus, such as feeding, sleep, and novelty seeking, emphasizing the close functional and developmental connection between these structures^13–16^. Despite the central roles of the hypothalamus and prethalamus in coordinating key bodily functions, the mechanisms governing their development remain poorly understood, particularly in relation to the specification of their diverse neuronal populations during embryonic development.

Single-cell RNA sequencing (scRNA-Seq) has proven effective in characterizing the cellular diversity of the adult hypothalamus^17–20^. Recent studies have extended these analyses to the developing hypothalamus and prethalamus in chick, mouse, and human models^21–26^. These approaches have provided detailed insights into the molecular identities of hypothalamic cell types and their developmental trajectories. However, the gene regulatory networks (GRNs) that control hypothalamic and prethalamic regionalization, neurogenesis, and cell type differentiation remain largely unknown. A major barrier to understanding these processes is the limited knowledge of how key developmental genes are regulated during hypothalamus and prethalamus formation. However, recent advances in single-cell multi-omics technologies, including the combination of scRNA-Seq with single-cell ATAC Sequencing (scATAC-Seq), now make it possible to link gene expression with chromatin accessibility and regulatory activity across development^27,28^. These approaches now allow a more precise dissection of the molecular mechanisms driving developmental processes, particularly when coupled with spatial transcriptomic analysis.

In this study, we present an integrated scRNA-Seq and scATAC-Seq analysis of the developing mouse hypothalamus and prethalamus, mapping the GRNs that control regionalization, neurogenesis, and cell type differentiation. Using these technologies, we identified key transcription factors (TFs) and regulatory mechanisms underlying hypothalamic and prethalamic development, including dynamic interactions guiding neuronal cell fate specification. Furthermore, we used this integrative approach to investigate the mechanisms of action of multiple TFs with broad and complex expression patterns in the developing hypothalamus and prethalamus, including *Isl1*, *Nkx2-2*, and *Dlx1/2*. Disrupting each of these genes leads to a loss of prethalamic identity and ectopic induction of genes specific to the supramammillary hypothalamus. In postnatal animals, selective loss of *Dlx1/2* in prethalamic and hypothalamic neuroepithelium leads to severe defects in the terminal differentiation of GABAergic neurons in the TRN and ZI that none, disrupting inhibition of excitatory thalamic neurons and resulting in hyperactivity in postnatal mice. These findings provide new insights into the regulatory architecture of the developing hypothalamus and highlight how disruptions in GRN function can alter neuronal circuitry and behavior.

## Results

### Developmental Dynamics of Hypothalamic and Prethalamic Cell Populations

To investigate the developmental trajectories and gene regulatory networks (GRNs) controlling hypothalamic patterning, neurogenesis, and cell fate specification, we generated integrated scRNA-Seq and scATAC-Seq datasets from the mouse hypothalamus and prethalamus. Key developmental time stages were sampled from embryonic day 11 (E11) to postnatal day 8 (P8), as described^24^. To analyze hypothalamic and prethalamic patterning and neurogenesis, we profiled the entire hypothalamus and prethalamus at E11, E12, E13, and E14, marking this period as the “Patterning and Neurogenesis Phase” (Fig 1A). Later stages of development, from E15, E16, E18, P4, and P8, excluding the telencephalon-derived preoptic area, constituted the “Differentiation Phase” (Fig 1A). In total, 340,000 cells were profiled using scRNA-Seq and 220,000 cells using scATAC-Seq (Table S1).

**Figure 1:**
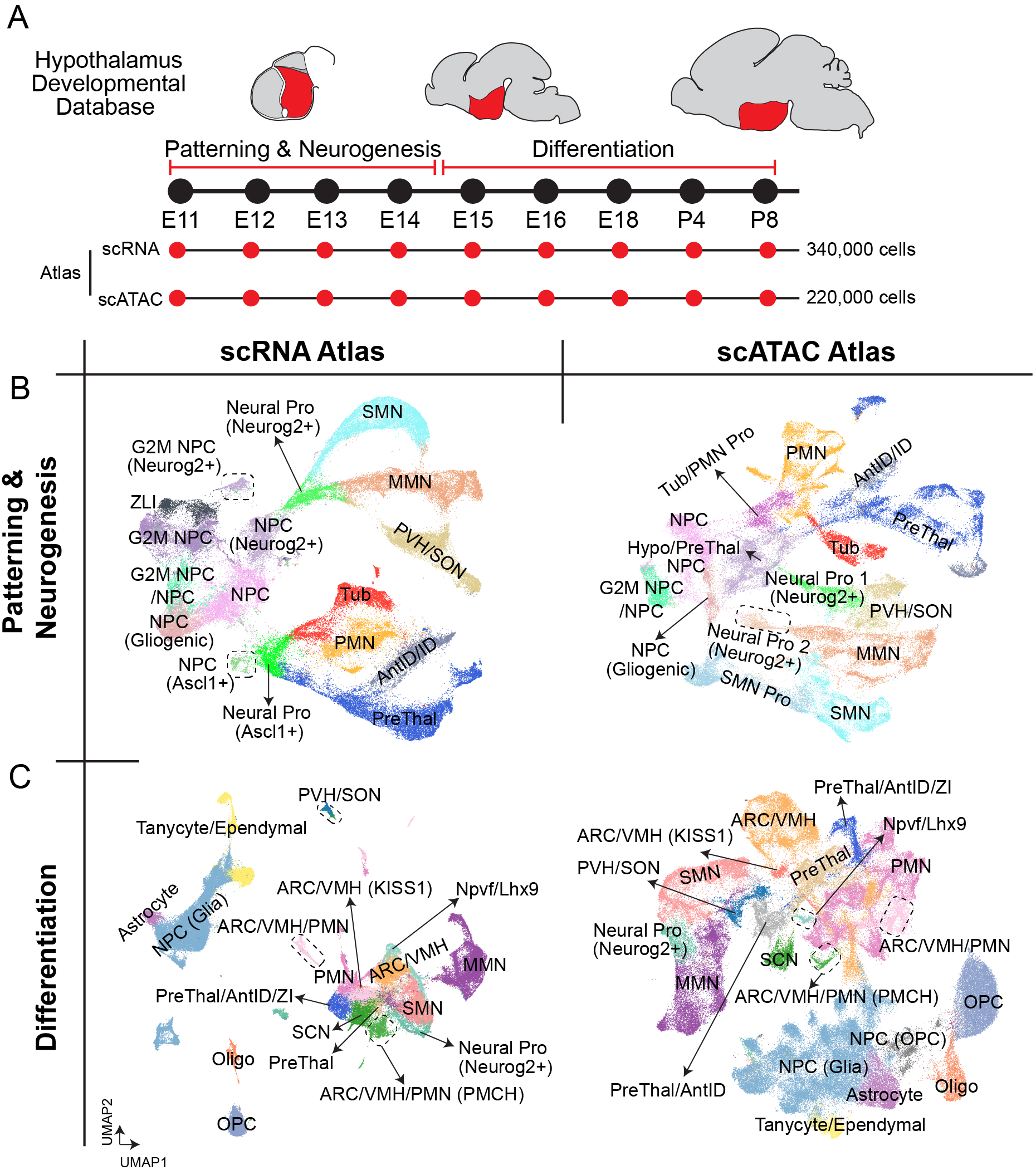
Single-cell atlas of mouse hypothalamic development using scRNA-Seq and scATAC-Seq (additional data in Fig S1, S2). **(A)** Schematic overview of hypothalamic development from embryonic day 11 (E11) to postnatal day 8 (P8). The timeline is divided into two developmental windows: *Patterning & Neurogenesis* (E11 to E14) and *Differentiation* (E15 to P8). Both scRNA-Seq and scATAC-Seq were performed across these stages to capture transcriptional and chromatin accessibility profiles across 340,000 and 220,000 cells, respectively. **(B)** UMAP projection of cell populations during the *Patterning & Neurogenesis* phase. Left: scRNA-Seq data showing clusters of neural progenitors and differentiating neuronal precursor cell types. Right: scATAC-Seq UMAP, highlighting chromatin accessibility states that correspond with transcriptional profiles observed in scRNA-Seq data. **(C)** UMAP projection of cell populations during the *Differentiation* phase. Left: scRNA-Seq data showing the emergence of mature cell types, including astrocytes, tanycytes, oligodendrocytes, and specific neuronal subtypes. Right: scATAC-Seq UMAP, highlighting chromatin accessibility states that correspond with cell type differentiation of hypothalamic neurons and glial cells.

During the Patterning and Neurogenesis phase, both scRNA-Seq and scATAC-Seq analyses revealed similar cell-type distributions, including distinct progenitor and early neuronal populations, consistent with previous findings^24^. G2/M-phase neural progenitors (NPCs, or primary progenitors) were distinguishable from G1/S-phase progenitors (Fig 1B, Tables S2, S3). A subset of NPCs, which increased in abundance over time, expressed high levels of glial markers (*Slc1a3, Aldh1l1*), suggesting these cells represent either fate-restricted gliogenic progenitors or quiescent radial glia (Fig 1B, Tables S2, S3). Furthermore, distinct NPCs expressing *Ascl1* or *Neurog2* were identified, potentially representing fate-restricted neurogenic progenitors (Fig 1B, Tables S2, S3). *Ascl1*-expressing neuronal progenitors (Neural Pro (Ascl1+)) give rise to neurons in the prethalamus, intrahypothalamic diagonal (ID), tuberal region, and premammillary hypothalamus (PMN) (Fig 1B). In contrast, *Neurog2*-expressing neuronal progenitors (Neural Pro (Neurog2+)), predominantly gave rise to neurons of the supramammillary nucleus (SMN), medial mammillary nucleus (MMN), and paraventricular nucleus of the hypothalamus (PVH)/supraoptic nucleus (SON) (Fig 1B). A subset of *Neurog2*-expressing tuberal progenitors was also observed (Fig 1B, S1A), corresponding to neurogenic progenitors in the ventromedial hypothalamus^29^. These two differentiation streams parallel findings from early hypothalamic development in chicks, where prethalamic-like and floorplate-like hypothalamic progenitors respectively give rise to GABAergic/tuberal and mammillary/supramammillary/paraventricular hypothalamic neurons^23^. The expression of transcription factors (TFs), and to a lesser extent the accessibility of their recognition motifs, delineated distinct spatial regions at this stage, further underscoring molecular mechanisms driving spatial and temporal specificity in hypothalamic development (Fig S1A).

Some differences between scRNA-Seq and scATAC-Seq profiles were observed at this stage. In the scRNA-Seq dataset, a well-defined cluster that selectively expresses markers (*Shh/Hes1*) of the zona limitans intrathalamica (ZLI), a transient structure separating the prethalamus from the sensory thalamus^30^, was evident but not distinguishable in the scATAC-Seq dataset (Fig 1B, Table S3). In contrast, while *Ascl1*- and *Neurog2*-expressing neurogenic progenitors were not readily further resolved into region-specific differentiation streams using scRNA-Seq data, region-specific identity was more evident in the scATAC-Seq data, especially within the *Neurog2* population, where region-specific neurogenic progenitor subtypes corresponding to SMN, MMN, and PVH were readily identified (Fig 1B).

As development progressed into E15-P8, the “Differentiation” phase when hypothalamic neurogenesis is essentially complete^30^, both transcriptional and chromatin accessibility landscapes reflected the emergence of mature hypothalamic cell types (Fig 1C). A small subpopulation of Neurog2-expressing neurogenic progenitors (Neural Pro (Neurog2+)) persisted at this stage (Fig 1C, Table S4), although Ascl1-expressing progenitors were no longer evident. Additional populations of committed cells, including various glial cell types, were also distinguishable by this point (Fig 1C, Tables S4, S5). GABAergic neuronal populations derived from precursors in the ID and prethalamus, including those in the suprachiasmatic nucleus (SCN) and the zona incerta (ZI), were also observed (Fig 1C, Tables S4, S5).

To better resolve specific hypothalamic cell types, we re-analyzed the scRNA-Seq datasets using higher clustering resolution (Fig S1B). This revealed distinct neuronal subpopulations corresponding to major hypothalamic neuronal subtypes, each defined by unique or overlapping neurotransmitter and neuropeptide expression patterns. Some neuronal subpopulations, such as *Pomc-* and *Agrp/Npy*-expressing neurons in the ARC, *Avp-*, and *Oxt*-expressing neurons in the PVH/SON, and *Hcrt-* and *Pmch-*expressing neurons in the lateral hypothalamus (LH), were already distinct at this stage (Fig S1B), suggesting earlier differentiation of these neurons compared to other hypothalamic cell types. Other less characterized neuronal subtypes, including cell clusters selectively expressing *Npw*^31,32^, *Grp/Cck*^33,34^, and *Npvf*^35^ in the dorsomedial hypothalamus (DMH), were also well-resolved (Fig S1B, Table S6). In the posterior hypothalamus, distinct subpopulations of SMN neurons (*Tac1, Calb2*) and MMN neurons (*Nts/Tac1/Cck, Npy/Cck, Cck*) were observed (Fig S1B, Table S6). The adult ZI structure was distinguished by selective expression of *Slc30a3*, as seen in both this scRNA-Seq dataset and *in situ* hybridization data from the Allen Brain Atlas (ABA) (Fig S1B).

To further explore the transcriptional diversity contributing to the specification of diverse neuronal subtypes at later stages of development, we re-analyzed scRNA-Seq data from the Patterning and Neurogenesis phase (E11-E14) using higher clustering resolution (Fig S2A, Table S7). Neuronal progenitors expressing *Ascl1* or *Neurog2* were subdivided into region-specific subpopulations based on high expression levels of TFs (Fig S2A, Table S7). *Bsx* expression was observed in a specific subcluster of cells within the PVH/SON progenitor population (Fig S2B). *In situ* hybridization data from the ABA confirmed that *Bsx* expression co-localized with the putative SON at E13.5, distinguishing the developing PVH and SON (Fig S2B). Further analysis of specific genes revealed distinct expression profiles for multiple neuropeptides and TFs (Fig S2C). The TFs *Sp9* and *Sp8* were selectively expressed in the prethalamus and AntID/ID progenitor clusters, respectively (Fig S2C). Neuropeptides such as *Cartpt* and *Ghrh* are localized to specific neuronal populations in the developing prethalamus and PVH/SON (Fig S2C). Similarly, *Pmch*, *Hcrt*, *Npvf,* and *Pnoc* already displayed restricted expression patterns at this stage (Fig S2C), indicating early maturation of these neuronal subtypes. *Npy* and *Sst* were also highly expressed in subsets of prethalamic neurons at this age, while *Gal* and *Prdm12* showed broad expression (Fig S2C).

### Gene Regulatory Networks Driving Hypothalamic and Prethalamic Progenitor Differentiation

To identify GRNs controlling hypothalamic regionalization and neurogenesis, we applied SCENIC+ analysis^27^, which integrates gene expression and chromatin accessibility to infer transcription factor activity. Using scRNA-Seq and scATAC-Seq datasets from early developmental time stages (E11-E14) (Fig 2A, S3, Table S8), we reconstructed GRNs that delineate hypothalamic regions and identified distinct networks specific to neuronal progenitors and neuronal precursors across most regions (Fig 2B, C, Fig S2A).

**Figure 2:**
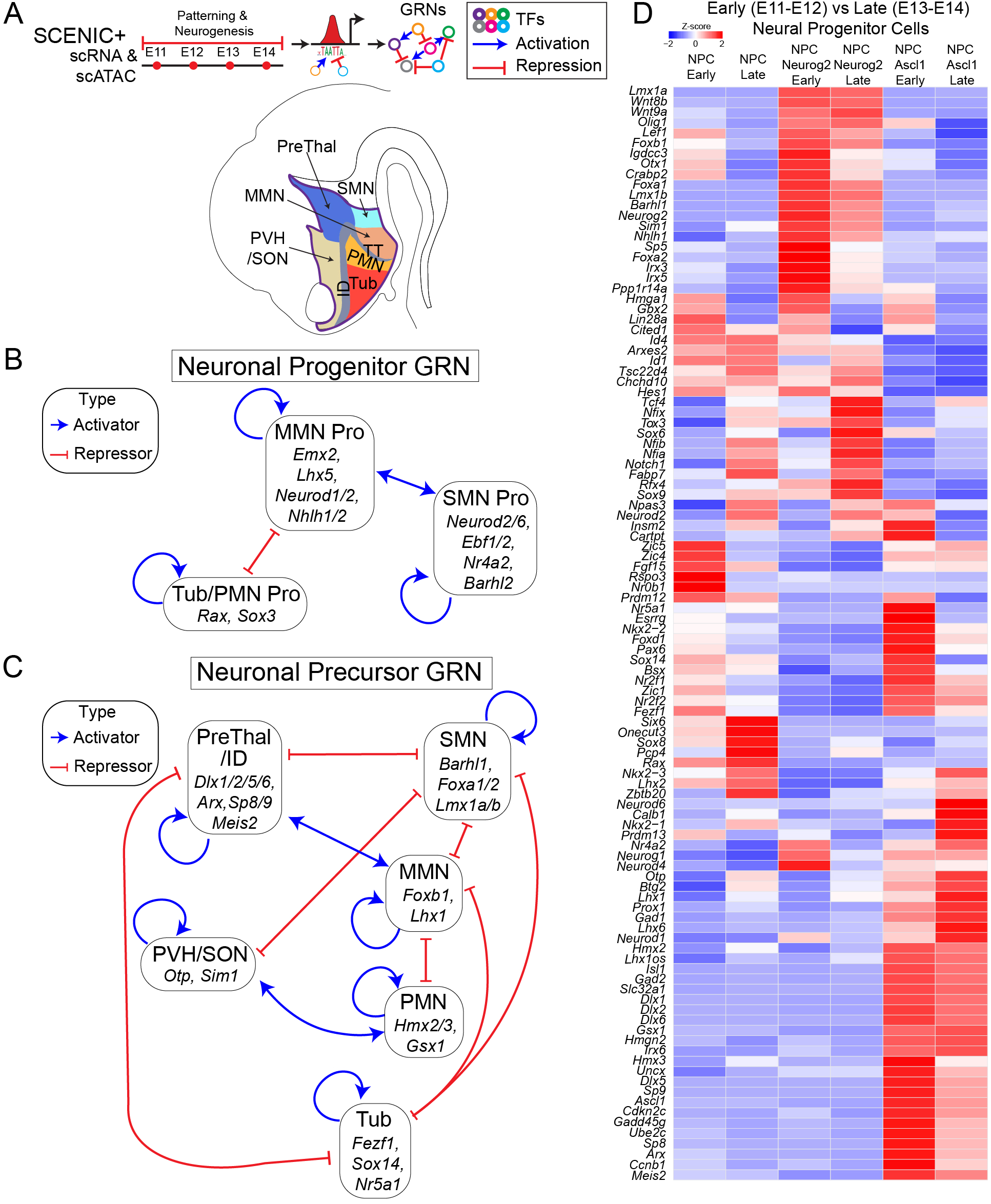
Gene regulatory networks controlling hypothalamic regionalization and temporal patterning (additional data in Fig S3, S4). **(A)** Top: Schematic of SCENIC+ analysis applied to scRNA-Seq and scATAC-Seq data from hypothalamic cells collected between E11 and E14. This analysis identifies gene regulatory networks (GRNs) composed of transcription factors (TFs) that act as activators (blue arrows) or repressors (red arrows). Bottom: Schematic illustration of the embryonic brain in the sagittal plane, with color-coded regions highlighting the major hypothalamic areas. **(B)** GRNs of Neuronal Progenitors in major hypothalamic regions (Fig S2A), including the tuberal (Tub), mammillary (MMN), premammillary (PMN), and suprachiasmatic (SMN). Key TFs regulating each region are shown, with arrows indicating activation (blue) or repression (red). **(C)** GRNs of Neuronal Precursors in major hypothalamic regions (Fig S2A), including the Tub, MMN, PMN, SMN, prethalamic/intrahypothalamic diagonal (PreThal/ID), and paraventricular/supraoptic (PVH/SON) nuclei. Key TFs regulating each region are shown, with arrows indicating activation (blue) or repression (red). **(D)** Heatmap illustrating differential expression of key TFs across early (E11–E12) and late (E13–E14) neural progenitor populations. Color intensity represents Z-scores of gene expression, with red indicating high expression levels and blue indicating low expression levels.

Within each region-specific GRN, TFs exhibited extensive cross-activation patterns, reinforcing their expression and that of other TFs in the same network. This positive feedback stabilizes cell identity and strengthens the robustness of region-specific gene expression programs (Fig 2B, C, Table S8). In parallel, most region-specific TFs repressed TFs from GRNs associated with other hypothalamic regions. For example, TFs active in the prethalamus/ID regions strongly inhibited TFs from GRNs promoting specification of the SMN and tuberal regions, and vice versa (Fig. 2C).

This cross-repression created a clear demarcation between hypothalamic regions, ensuring that progenitor cells committed to specific regional identities. Such interactions likely preserve distinct molecular and functional domains throughout hypothalamic development. These regulatory relationships (Fig 2C) closely match regulatory relationships seen at much earlier stages of chick hypothalamic regionalization^23^. These cross-repressive regulatory relationships likely ensure the robust selection of discrete states and maintain the distinct identity of spatially and functionally divergent regions from early hypothalamic regionalization through the end of neurogenesis at E14.

Interestingly, the TFs in the SMN and MMN neuronal progenitor networks frequently cross-activated one another during early development (Fig 2B). However, this pattern was not observed for the prethalamus/ID, tuberal, or PMN regions. In later-stage neuronal precursors, we observed extensive cross-inhibitory interactions between the GRNs active in the SMN, MMN, and PVH/SON regions, indicating the tightening of regulatory boundaries as these regions matured (Fig 2C). In contrast, no significant cross-inhibitory regulation was detected between the tuberal and PMN regions (Fig 2C). Instances of common TFs between GRNs controlling specification of non-contiguous regions, such as the PVH/SON and PMN, or the prethalamus/ID and MMN (Fig S3), suggest common regulatory elements influencing their development despite their spatial separation.

In all regions of the central nervous system (CNS), progenitors exhibit temporal patterning, where both the timing of neurogenesis and the competence to generate specific cell fates change dynamically over time^36,37^. This analysis identified candidate regulators of temporal identity within hypothalamic progenitors, revealing distinct gene expression profiles in early-stage (E11-E12) versus late-stage (E13-E14) progenitors. A small subset of genes was enriched either in the early or late stages of both primary NPCs and neurogenic progenitors, including *Neurog2*- and *Ascl1*-expressing cells (Fig 2D, Table S9). Genes enriched in both types of early-stage progenitors included *Hmga1, Lin28a*, *Cited1*, and *Gbx2*, while late-stage progenitors showed elevated expression of *Tcf4, Npas3, Nfia/b/x, Zbtb20,* and *Tox3* (Fig 2D). Many of these genes are implicated in temporal patterning across other CNS regions, such as the cerebral cortex, retina, and spinal cord^28,38–41^, implying that common mechanisms control temporal regulation of neurogenesis in multiple CNS structures.

Most stage-specific genes, however, showed dynamic expression only in particular progenitor classes. For example, early-stage primary NPCs showed selective enrichment of *Zic4*, *Zic5*, *Fgf15*, *Rspo3*, and *Nr0b1*, while *Six6*, *Onecut3*, and *Rax* were more prominently expressed in late-stage NPCs (Fig 2D). In neurogenic progenitors, *Neurog2+* progenitors exhibited early-stage expression of SMN-specific factors such as *Lmx1a*, *Foxa1/2*, and *Irx3/5*, while *Ascl1+* progenitors showed early-stage enriched expression of prethalamus-specific genes, including *Dlx1/2/6*, *Sp8/9*, and *Arx* (Fig 2D). These temporally dynamic TFs play crucial roles in specifying SMN and prethalamic identities, respectively.

We also observed significant differences in the temporal dynamics of Notch pathway genes. *Hes1* was strongly enriched in early-stage NPCs and *Neurog2+* progenitors, while *Notch1* expression increased in late-stage progenitors (Fig 2D). However, these changes were not observed in *Ascl1+* progenitors, where Notch pathway gene expression remained low and relatively constant over time. Similarly, NFI family TFs, which are key regulators of late-stage temporal identity in the retina^28,38^, were selectively active in late-stage NPCs and *Neurog2+* progenitors, but absent in *Ascl1+* progenitors at E14.

### A Widespread Role for Transcription Factors Controlling SMN Development in Susceptibility to Metabolic and Neuropsychiatric Disease

To explore potential links between hypothalamic development and disease-related genetic traits, we integrated GRN analysis with genome-wide association study (GWAS) data from the EMBI-EBI database^42^. This linked transcription factor activity in hypothalamic progenitor populations to genetic variants associated with complex traits and diseases, providing insights into how early transcription regulation in specific cell types in the developing hypothalamus and prethalamus might influence disease susceptibility (Fig S4). We observed several patterns. First, TFs broadly active in progenitor cell populations, such as NSC/NPC and *Neurog2*+ progenitors, showed strong associations with a broad range of neurological and metabolic traits such as kidney disease, headache, and diabetes. Second, TFs broadly active in hypothalamic neural precursor cells were linked to Alzheimer’s disease and hypertension. Third, TFs selectively active in tuberal and PMN progenitors were associated with traits including response to antidepressants and Lewy body disease, suggesting a selective role of these progenitors in neuropsychiatric health^45–47^. Fourth, TFs broadly active both in multiple progenitor subtypes in SMN precursors, but not in other hypothalamic neuronal precursor subtypes, were linked to metabolic traits such as body mass, fat distribution, and TSH levels, as well as to behavioral traits such as alcohol consumption, smoking, and depression. This was also the case for TFs that were selectively active in the SMN precursors, which were linked traits including BMI, bipolar disorder, schizophrenia, cognition, IQ, and smoking initiation, hinting at an additional unexpected role for the SMN in both metabolic and psychiatric health. This highlights a potentially uncharacterized role of the SMN in metabolic regulation and motivated behavior ^43,44^. Overall, these findings suggest that TFs associated with hypothalamic progenitors, and the SMN in particular, may be pivotal in establishing developmental trajectories that influence susceptibility to a range of metabolic, neuropsychiatric, and cognitive disorders^48,49^.

### Gene Regulatory Networks Controlling Hypothalamic Neurotransmitter and Neuropeptide Expression

Each hypothalamic region contains numerous molecularly and functionally distinct neuronal cell types, often characterized by the selective expression of neurotransmitters and neuropeptides^20,24,26^. To investigate GRNs controlling neurotransmitter and neuropeptide identity, we applied SCENIC+ analysis to scRNA-Seq and scATAC-Seq data obtained from neurons collected between E15 and P8 (Fig 3A, S5, Table S10). This approach enabled us to map TFs regulating key markers of mature neuronal identity, providing insights into how hypothalamic regionalization impacts neurotransmitter and neuropeptide expression concurrently.

**Figure 3:**
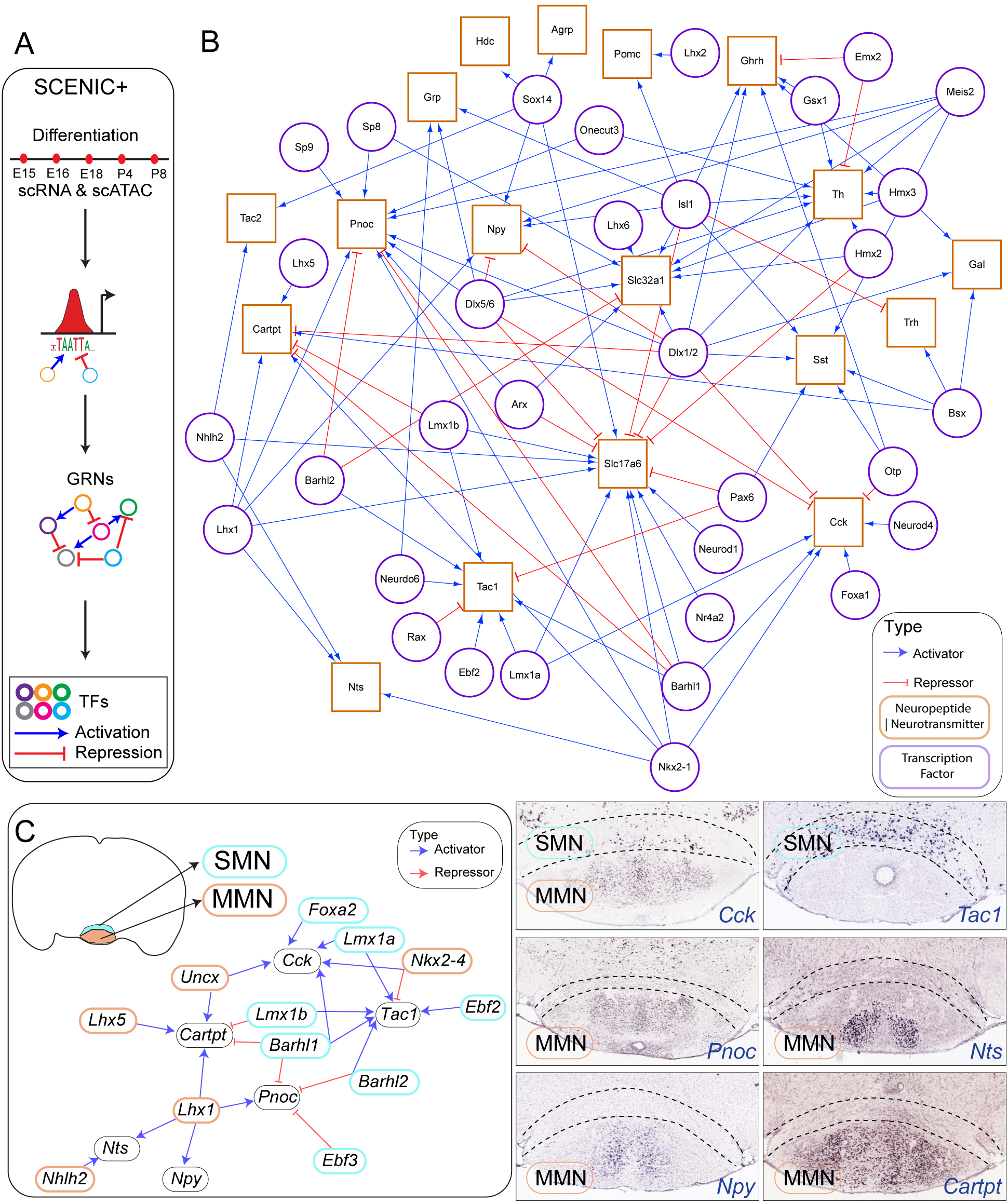
Gene regulatory networks controlling hypothalamic neurotransmitter and neuropeptide expression (additional data in Fig S5). **(A)** Schematic of SCENIC+ analysis applied to the *Differentiation* phase (E15 to P8), using scRNA-Seq and scATAC-Seq data to construct gene regulatory networks (GRNs). Transcription factors (TFs) act as activators (blue arrows) or repressors (red arrows) to regulate neuropeptide and neurotransmitter genes. **(B)** GRN highlighting transcription factor interactions and neuropeptide/neurotransmitter-encoding genes across hypothalamic regions. Activating (blue) and repressing (red) interactions are shown between TFs (purple) and neuropeptide/neurotransmitter genes (yellow). **(C)** Left: regional GRNs specific to the supramammillary nucleus (SMN) and mammillary nucleus (MMN), showing key TFs and neuropeptides/neurotransmitters. Right: validation images from the Allen Brain Atlas (right panels) display *in situ* hybridization images for *Tac1*, *Cck*, *Pnoc*, *Npy*, *Nts*, and *Cartpt*, highlighting their expression in SMN and MMN.

Many TFs involved in the early stages of hypothalamic regionalization continue to regulate the expression of neurotransmitter-related genes in a feed-forward manner. For example, prethalamic/ID regulators such as *Dlx1/2/5/6*, *Arx*, *Sp8*, and *Meis2* directly activate genes involved in GABAergic neurotransmission, such as *Slc32a1* (Fig 3B). These same GABAergic genes are directly repressed by TFs specific to Neurog2+ glutamatergic neurons, including *Barhl2*, highlighting the antagonistic regulation between GRNs specifying excitatory and inhibitory neurons. Conversely, genes mediating glutamatergic neurotransmission, such as the vesicular glutamate transporter *Slc17a6*, are activated by both *Neurog2*-specific TFs (*Barhl1, Nhlh2, Nr4a2, Lmx1a, Neurod1*) and tuberal progenitor-specific factors (*Nkx2-1, Sox14*). These glutamatergic genes are repressed by prethalamic-specific TFs (Fig 3B). Not all activators of GABAergic genes are confined to prethalamic regions, however. TFs such as *Hmx2/3* and *Isl1*, expressed across the tuberal, premammillary, and prethalamic regions (Fig 3B), also regulate GABAergic gene expression^50,51^.

Neuropeptides and monoamines exhibit complex and variable patterns of regulation by TFs that specify regional identity. Several known regulatory relationships were recapitulated in this analysis (Fig 3B). For instance, consistent with previous studies, *Otp* activates *Sst* expression, while *Npy* expression is activated by *Isl1* and repressed by *Dlx1/2*^50,52^. Beyond validating known interactions, this uncovered novel regulatory patterns in which neuropeptide expression is controlled by mutually antagonistic GRNs in region-specific contexts, particularly in the SMN and MMN.

In the SMN *Tac1*, which encodes substance P, is selectively expressed in specific subpopulations of neurons, but is absent in the MMN (Fig 3C). In contrast, *Cck* is strongly expressed in specific subsets of both SMN and MMN neurons. Moreover, neuropeptides such as *Pnoc*, *Npy, Nts*, and *Cartpt* are exclusively expressed in MMN neurons, with no expression in SMN neurons (Fig 3C). GRN analysis revealed that *Tac1* expression is directly activated by SMN-specific TFs, including *Lmx1a/b*, *Foxa2, Barhl1/2,* and *Ebf2*, while it is repressed by the MMN-specific factor *Nkx2-4* (Fig 3C). In contrast, *Cck* expression is activated by both SMN- and MMN-specific factors, such as *Nkx2-4* and *Uncx*, indicating a shared regulatory mechanism for this neuropeptide across different regions. Conversely, MMN-specific neuropeptides (*Pnoc, Npy, Nts, Cartpt*) are activated by MMN-specific TFs, including *Lhx1/5, Uncx,* and *Nhlh2*, but are directly repressed by SMN-specific TFs, reinforcing the mutual antagonism between GRNs governing neurotransmitter and neuropeptide expression in distinct hypothalamic regions (Fig 3C).

### Identification of Dopaminergic Neurons in the Hypothalamus and Their Regulatory Networks

Hypothalamic dopamine signaling plays a critical role in modulating numerous physiological processes, including reward, feeding, reproductive behavior, and neuroendocrine regulation^53^. Given its importance, we next analyzed *Th*-expressing dopaminergic neurons across the hypothalamus to identify the transcriptional networks governing their differentiation. We identified several *Th*-expressing clusters throughout the hypothalamus, including cells located in the prethalamus, which likely give rise to A13 dopaminergic neurons located in the ZI (Fig S6A). Additionally, *Th*-expressing clusters were found in the PMN/Tuberal regions, PVH/SON, and SMN (Fig S6A, Table S11), potentially corresponding to A12, A15 and A11 subtypes respectively. To investigate the regulatory mechanisms controlling *Th* expression, we performed GRN analysis, which identified a set of TFs critical for dopamine expression. In presumptive A13 precursors, these included *Dlx1/2/5/6,* and *Meis2*, which are also predicted to be crucial for specifying both prethalamic identity and GABAergic neurons (Fig S6B). In immature A12 neurons, both PMN-enriched TFs such as *Hmx2/3* directly activate *Th* along with more broadly expressed TFs such as *Isl1, Pbx3,* and *Mef2c* (Fig S6B). Furthermore, MMN-enriched TFs *Emx2* and *Nkx2-4* were found to act as direct repressors of *Th* expression, thereby potentially restricting the specification of both A11 and A12 neurons (Fig S6B).

We also observed the co-expression of additional neuropeptides, TFs, and receptors across the *Th*-expressing clusters, which revealed additional layers of functional diversity (Fig S6C). For example, *Arx*, a marker of GABAergic neurons^54,55^, and *Esr1*, a key regulator of hormonal feedback^56,57^, were expressed in presumptive A13 and A12 clusters, respectively. Moreover, neuropeptides such as *Trh* and *Tac1* were detected in distinct *Th*-expressing clusters, highlighting the heterogeneity within hypothalamic dopaminergic neurons (Fig S6C).

### Genetic Analysis Reveals that the LIM Homeodomain Factors Lhx1 and Isl1 Control Regional Identity in Developing Hypothalamus and Prethalamus

GRN analysis identified key roles for multiple LIM homeodomain TFs that are expressed in complex, overlapping patterns during early hypothalamic development. One such TF is *Lhx1*, which is expressed in the prethalamus and ID, as well as the developing mammillary region, encompassing both *Ascl1*-expressing prethalamic progenitors and *Neurog2*-expressing mammillary progenitors^24,30^. A second LIM homeodomain *Isl1*, which is expressed in both neurogenic progenitors and postmitotic precursors across a broad, contiguous domain extending ventrally from the prethalamus, through the ID, and into the tuberal and PMN regions. This expression pattern does not strictly correspond to any identified anatomical domains^30^. To validate these predicted regulatory relationships, we selectively disrupted *Lhx1* and *Isl1* in early hypothalamic and prethalamic neuroepithelial cells from E9.5 onwards using the *Foxd1*-*Cre* transgenic line^58^. Homozygous mutants and controls at E12.5 were profiled using scRNA-Seq, with scATAC-Seq also performed for *Isl1* mutants.

*Lhx1,* which was predicted to activate a subset of TFs specific to the prethalamus, ID, and MMN while repressing TFs specific to the PMN and SMN (Fig 4A). *Lhx1* mutant showed a modest reduction in cells expressing MMN-specific markers, accompanied by a modest increase in genes enriched in PMN progenitors such as *Lhx2* and *Tcf7l2* (Fig 4B). These findings indicate that while *Lhx1* is essential for the full specification of MMN identity, its disruption does not eliminate MMN progenitors. Interestingly, neuronal subpopulations that were essentially absent in controls appeared predominantly in *Lhx1* mutants (Fig 4C, Table S12). These mixed populations co-expressed markers of SMN and MMN progenitors, which normally exclusively give rise to glutamatergic neurons, and GABAergic markers such as *Gad1* (Fig 4D, S7, Table S13). This suggests that in the absence of *Lhx1*, a subset of hypothalamic progenitors adopt a mixed identity, expressing both SMN/MMN and GABAergic markers.

**Figure 4:**
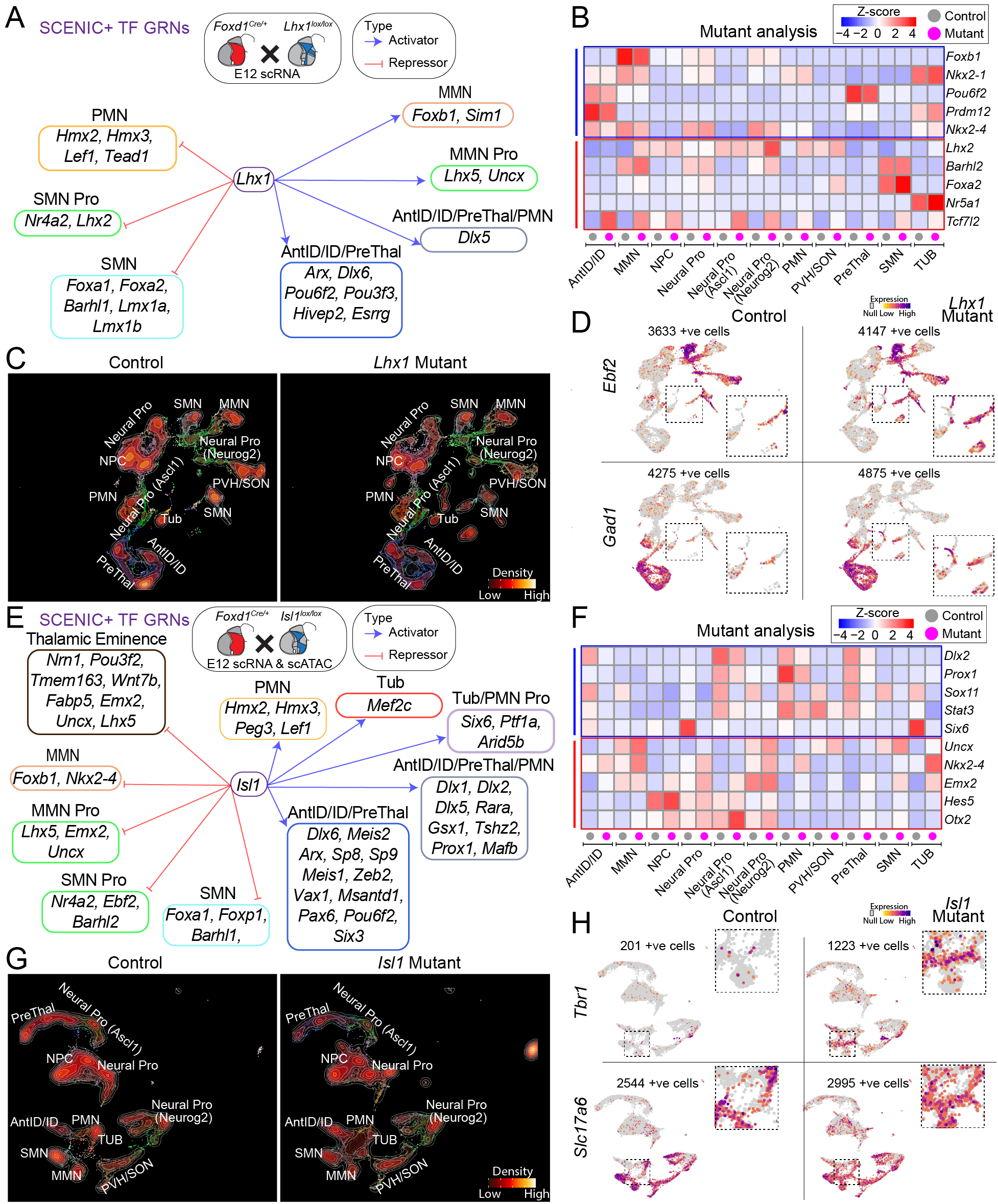
Disruption of hypothalamic GRNs in *Lhx1* and *Isl1* mutant mice (additional data in Fig S7, S8). **(A)** Gene regulatory networks (GRNs) regulated by *Lhx1* in hypothalamic cells using SCENIC+ analysis. **(B)** Heatmap showing transcriptional changes in *Lhx1* mutant, with color intensity indicating differential gene expression levels. **(C)** UMAP plots comparing control and *Lhx1* mutant hypothalamus. Brighter colors represent higher cell density. **(D)** Expression of *Ebf2* and *Gad1* in control versus *Lhx1* mutant hypothalamus. **(E)** GRNs regulated by *Isl1* in hypothalamic cells using SCENIC+ analysis. **(F)** Heatmap showing transcriptional changes across multiple hypothalamic progenitor populations in *Isl1* mutant, with color intensity indicating differential gene expression levels. **(G)** UMAP plots comparing cell populations in control and *Isl1* mutant hypothalamus. Brighter colors represent higher cell density. **(H)** Expression of *Tbr1* and *Slc17a6* in control versus *Isl1* mutant hypothalamus.

GRN analysis predicted that *Isl1* directly activates TFs specific to the prethalamus, ID, PMN, and tuberal regions while repressing TFs associated with the SMN, MMN, and thalamic eminence (EmThal) (Fig 4E, F, Fig S8, Table S14, S15). The EmThal, a domain in the anterior prethalamus that predominantly gives rise to excitatory neurons^59,60^. Analysis of *Isl1* mutants confirmed these initial predictions, with the most dramatic changes observed in the prethalamus and PMN regions of *Isl1* mutants. The relative fraction of both prethalamus and PMN cells was substantially reduced in *Isl1* mutants compared to the controls (Fig 4G, Table S16). Additionally, we observed a mutant-specific cell cluster that co-expressed early EmThal-like markers, including the transcription factor *Tbr1* and glutamatergic markers such as *Slc17a6* (Fig 4H)^61^, along with SMN and MMN markers (Fig S8A).

### *Nkx2.2* Promotes Prethalamic Specification and Represses SMN and MMN Identity

Previous studies on early chick hypothalamic development demonstrated that antagonistic signaling between prethalamic and floorplate-like neuroepithelial cells initiates hypothalamic regionalization^23^. Prethalamic-like cells subsequently give rise to *Ascl1*-expressing prethalamic and tuberal regions, while floorplate-like cells generate *Neurog2*-expressing SMN and MMN regions^23,24^. GRN analysis suggests that these regulatory relationships are conserved in mammals. Specifically, TFs promoting prethalamic and tuberal specification repress the differentiation of SMN and MMN regions, and vice versa.

Building on our initial observations of *Isl1* and *Lhx1* mutants, we explored the role of additional TFs predicted to regulate hypothalamic regionalization. We focused on *Nkx2-2*, a known downstream target of Shh signaling^62^, expressed in the dorsal tuberal region, ventral prethalamus, and ID, and demarcating the separation of alar and basal hypothalamic regions^30^. GRN analysis predicted that Nkx2-2 directly activates a subset of TFs specific to the tuberal hypothalamus and prethalamus/ID regions, while selectively repressing genes specific to the SMN (Fig 5A). This suggests that *Nkx2-2* maintains regional boundaries by promoting prethalamic and tuberal fates while repressing caudal hypothalamic regions, such as the SMN.

**Figure 5:**
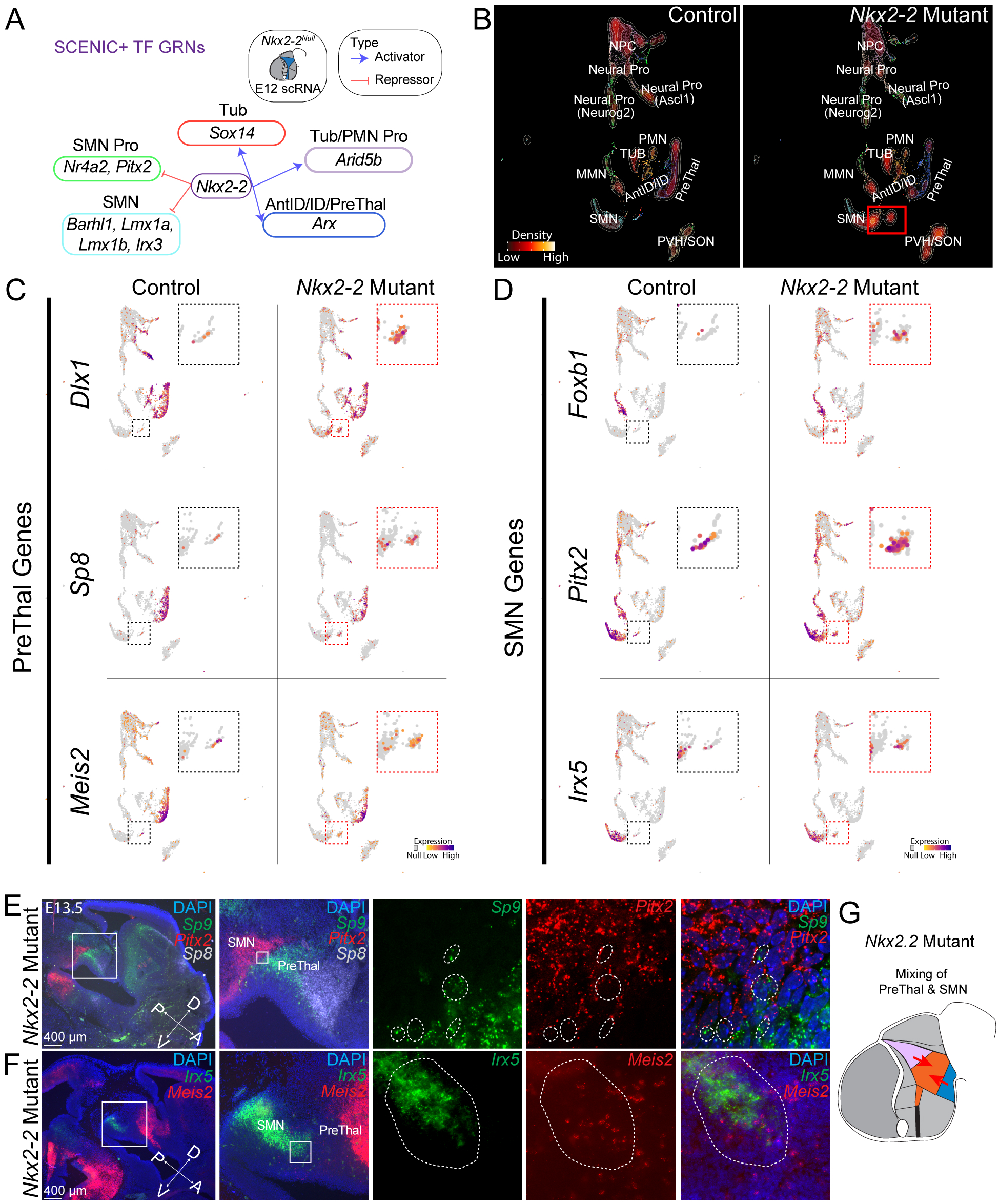
Disruption of *Nkx2.2* leads to the generation of cells with mixed prethalamic and SMN identity (additional data in Fig S9). **(A)** Gene regulatory networks (GRNs) showing *Nkx2.2* as a repressor of supramammillary nucleus (SMN)-promoting transcription factors and an activator of prethalamic (PreThal) identity. **(B)** UMAP projections of control and *Nkx2.2* mutant hypothalamus, showing a reduction in prethalamus populations and the expansion of SMN-like cells in the mutant. Brighter colors represent higher cell density. **(C)** Feature plots of key PreThal genes (*Dlx1*, *Sp8*, *Meis2*) in control and *Nkx2.2* mutant hypothalamus, showing reduced expression in mutant animals. **(D)** Feature plots of SMN-specific genes (*Foxb1*, *Pitx2*, *Irx5*) in control and *Nkx2.2* mutant hypothalamus, showing ectopic expression in the prethalamic region of mutant animals. **(E)** *In situ* hybridization in *Nkx2.2* mutant hypothalamus showing overlapping expression of *Sp8* and *Pitx2* in the prethalamic region, whereas *Sp8* is typically restricted to the prethalamus and *Pitx2* to the SMN in controls. Scale bar: 400 μm. **(F)** *In situ* hybridization in *Nkx2.2* mutant hypothalamus showing co-expression of *Irx5* (SMN marker) and *Meis2* (prethalamus marker) in the prethalamus of *Nkx2.2*-mutant hypothalamus, validating the mixing of neuronal identities. **(G)** Schematic summarizing the disruption of prethalamic identity and the resulting generation of cells with mixed prethalamic and SMN identity in *Nkx2.2* mutants.

To test these predictions, we generated homozygous *Nkx2-2* mutants and profiled the hypothalamus and prethalamus using scRNA-Seq. Significant changes in regional identity were observed in the *Nkx2-2* mutants. As predicted, the relative number of prethalamus/ID neurons was reduced, while increased numbers of SMN neurons were observed (Fig 5B, Table S17). Interestingly, a novel cell cluster emerged in the mutants that co-expressed markers from both the prethalamus and SMN/MMN regions (Fig 5C, D, Fig S9, Table S18). Reduced expression of prethalamic genes and ectopic expression of SMN/MMN progenitor genes such as *Pitx2* were also observed in the prethalamus cluster (Fig 5C, D, Fig S9, Table S18).

Fluorescent *in situ* hybridization confirmed the presence of these mixed-identity cells in the posterior prethalamic region (Fig 5E, F). Specifically, this region in *Nkx2-2* mutants co-expressed the prethalamus marker *Sp9* together with the SMN/MMN marker *Pitx2* (Fig 5E). Similarly, co-expression of the prethalamus marker *Meis2* and the SMN marker *Irx5* was observed in the prethalamus of *Nkx2-2* mutants (Fig 5F). These results indicate that loss of function of *Nkx2-2* results in a disruption of GRNs that normally segregate the prethalamus/ID and SMN/MMN regions, leading to the generation of cells with mixed identities (Fig 5G).

### *Dlx1/2* Promote Prethalamic Identity and GABAergic Differentiation While Repressing SMN Identity

Finally, we examined this analysis to *Dlx1/2*, which are well-characterized as key regulators of telencephalic GABAergic neuronal differentiation^63,64^. Our analysis indicates that, within the hypothalamus, *Dlx1/2* activate the expression of *Ascl1* and multiple TFs predicted to promote prethalamus, ID, and PMN identities (Fig 6A). Concurrently, *Dlx1/2* repress *Neurog2* and other genes involved in the specification of SMN, MMN, and a subset of tuberal cell types (Fig 6A). Overall, *Dlx1/2* are predicted to be among the strongest regulators of regional identity and neurogenesis in the developing hypothalamus and prethalamus, as well as orchestrating the balance between GABAergic and glutamatergic neuronal fates (Fig 3B). To further explore the role of *Dlx1/2* in hypothalamic and prethalamus regionalization, we generated conditional mutants for *Dlx1/2* using the *Foxd1-Cre* transgenic line and performed scRNA-Seq and scATAC-Seq analyses to examine the impacts of *Dlx1/2* deletion on transcriptional and chromatin landscapes (Fig 6B, S10, Table S19).

**Figure 6:**
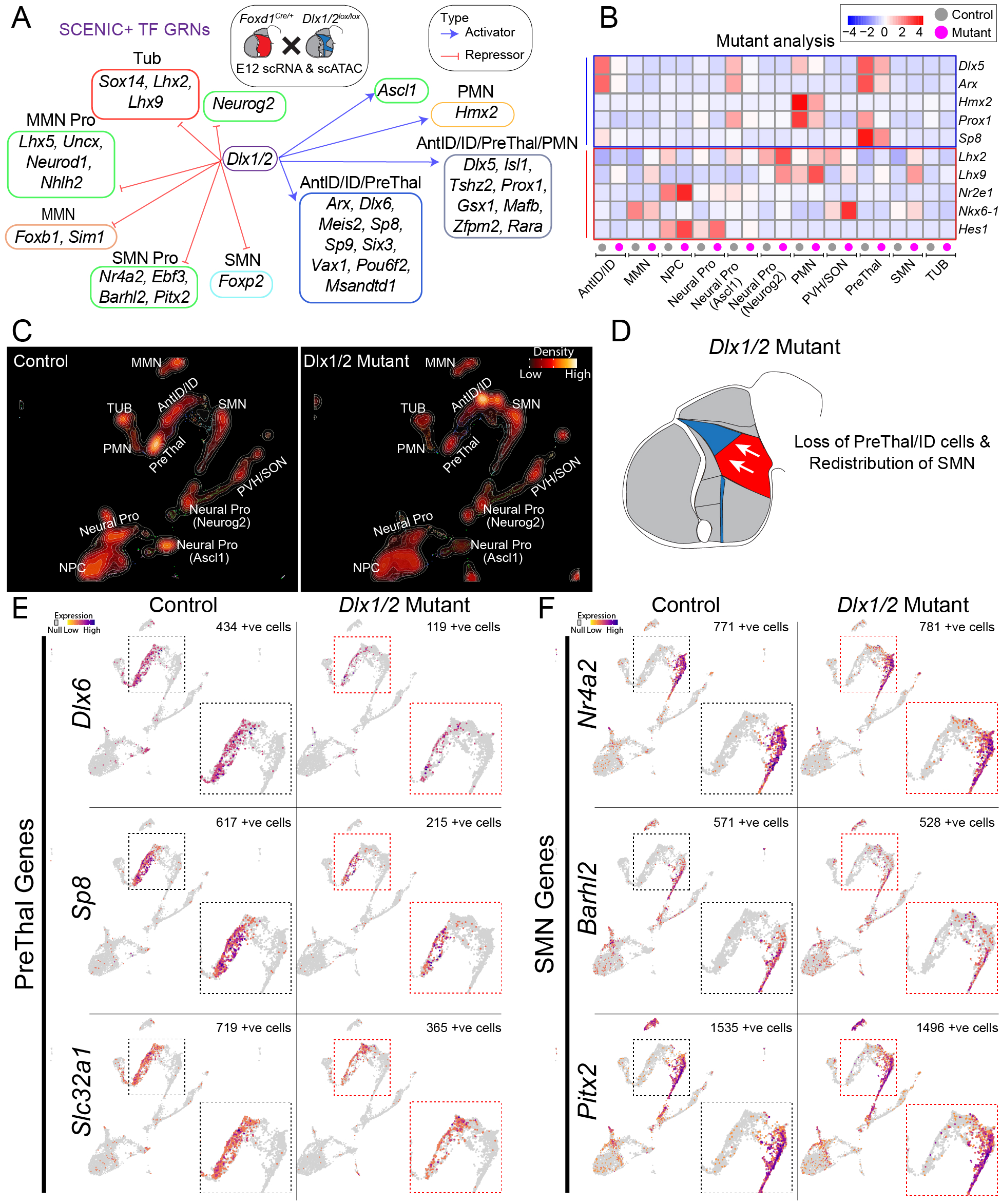
Disruption of *Dlx1/2* leads to loss of prethalamic identity and expansion of SMN markers (additional data in Fig S10, S11). **(A)** Gene regulatory networks (GRNs) showing *Dlx1/2* as activators of prethalamic (PreThal) transcription factors and repressors of SMN markers. **(B)** Heatmap showing transcriptional changes in *Dlx1/2* mutants compared to controls, with color intensity indicating differential gene expression. **(C)** UMAP projections comparing control and *Dlx1/2* mutant hypothalamus, displaying reduced prethalamus and intrahypothalamic diagonal (ID) populations in *Dlx1/2* mutants, along with an expansion of SMN neurons. Brighter colors represent higher cell density. **(D)** Schematic summarizing the loss of prethalamus and ID identity and the redistribution of SMN-like cells in *Dlx1/2* mutants. **(E)** Feature plots of key prethalamic genes (*Dlx6*, *Sp8*, *Slc32a1*) in control and *Dlx1/2* mutants, showing downregulation of these genes in mutants. **(F)** Feature plots of SMN-specific genes (*Nr4a2*, *Barhl2*, *Pitx2*) in control and *Dlx1/2* mutants, showing ectopic upregulation of these genes in the mutant prethalamus and ID.

As predicted by GRN analysis, *Dlx1/2* mutants exhibited a dramatic reduction in *Ascl1*-expressing progenitors, along with a significant loss of prethalamus/ID and PMN neural precursors (Fig 6B-D, Table S20, S21). GABAergic markers, including *Slc32a1,* were markedly reduced in the mutants, consistent with the loss of inhibitory neuronal identity (Fig 6E). This reduction highlights the critical role of *Dlx1/2* in activating transcriptional programs that drive GABAergic neurogenesis in the prethalamus and ID. In addition, *Dlx1/2* mutants exhibited an expansion of SMN-like cells, with genes typically restricted to the SMN, such as *Nr4a2*, and *Barhl2*, being ectopically expressed in clusters that would normally give rise to prethalamic neurons (Fig 6F).

Histological analysis at E13.5 confirmed these findings, showing a widespread reduction in the expression of *Dlx2, Isl1, Arx, Dlx5, Gad2, Hmx2, Meis2, Sp8, Sp9,* and *Pax6* in *Dlx1/2* mutants (Fig S11). Alongside this loss of prethalamus and ID identity, there was a dorsal expansion of cells expressing SMN markers, such as *Foxa1* and *Pitx2* (Fig S11B, C), consistent with the scRNA-Seq data (Fig 5F). These results imply that in the absence of *Dlx1/2*, prethalamic and ID progenitors lose expression of region-specific markers and undergo partial conversion to SMN-like identity.

To assess postnatal effects of *Dlx1/2* mutants, we first performed scRNA-Seq on the hypothalamus and the prethalamus-derived ZI and ventral TRN at P8 (Fig S12A). Persistent changes in the prethalamus were observed (Table S22), including a notable loss of specific neuronal subpopulations in the ARC, such as *Agrp* and *Ghrh*-expressing neurons, consistent with previous findings^52^. Additionally, there was a reduction in *Sst, Th*, and *Pnoc*-expressing neurons, as predicted by earlier developmental studies (Fig 3B, Table S22). Interestingly, there was a dramatic downregulation of *Npw* (Fig S12B), which is transiently expressed during development^32^ and co-expressed with *Th* in the postnatal DMH (Fig S6C).

Despite the severity of the early developmental phenotype (Fig 6), *Foxd1-Cre;Dlx1/2^lox/lox^* mice (*Dlx1/2* mutants) are viable and born at expected Mendelian ratios. *Foxd1-Cre;Dlx1/2^lox/lox^* mice were efficiently generated by crossing homozygous *Dlx1/2^lox/lox^* mice with *Foxd1-Cre;Dlx1/2^lox/+^*mice. This cross resulted in ∼12.5% of the offspring being heterozygous for *Foxd1-Cre* and homozygous for *Dlx1/2^lox/lox^*. However, *Dlx1/2* mutants displayed significant reductions in body weight and body length (Fig S12C, Table S23), consistent with previous findings in the tuberal hypothalamus-specific *Dlx1/2* mutants^52^. These growth deficits likely relate to disruptions in *Ghrh*-expressing neurons and an altered balance of *Agrp*-expressing neurons in the ARC^52^. *Dlx1/2* mutants showed poor growth on a standard chow diet, with nearly all individuals dying within the first post-weaning week (Fig S12D, Table S24). However, survival significantly improved when a gel-based food was provided *ad libitum* (Fig S12D, Table S25). The mechanism underlying the failure to thrive on standard chow remains unclear.

Given the substantial loss of neurons in prethalamus-derived structures in *Dlx1/2* mutant embryos, we specifically focused on the TRN and ZI in post-weaned *Dlx1/2* mutants. A reduction in the number of neurons was also evident in these regions (Fig 7A). We performed snRNA-Seq on the TRN and ZI, along with adjacent thalamic regions at P21 (Fig S13A, B, Tables S26, S27), and Xenium spatial transcriptomics, which was performed using a custom panel of genes selectively expressed in individual hypothalamic and prethalamic cell types (Table S28). Both analyses revealed a marked reduction in the number of both *Pvalb*/*Meis2* and *Sst/Penk*-expressing neurons within the ZI and TRN (Fig 7B, S13C, Tables S26, S27, S29).

**Figure 7:**
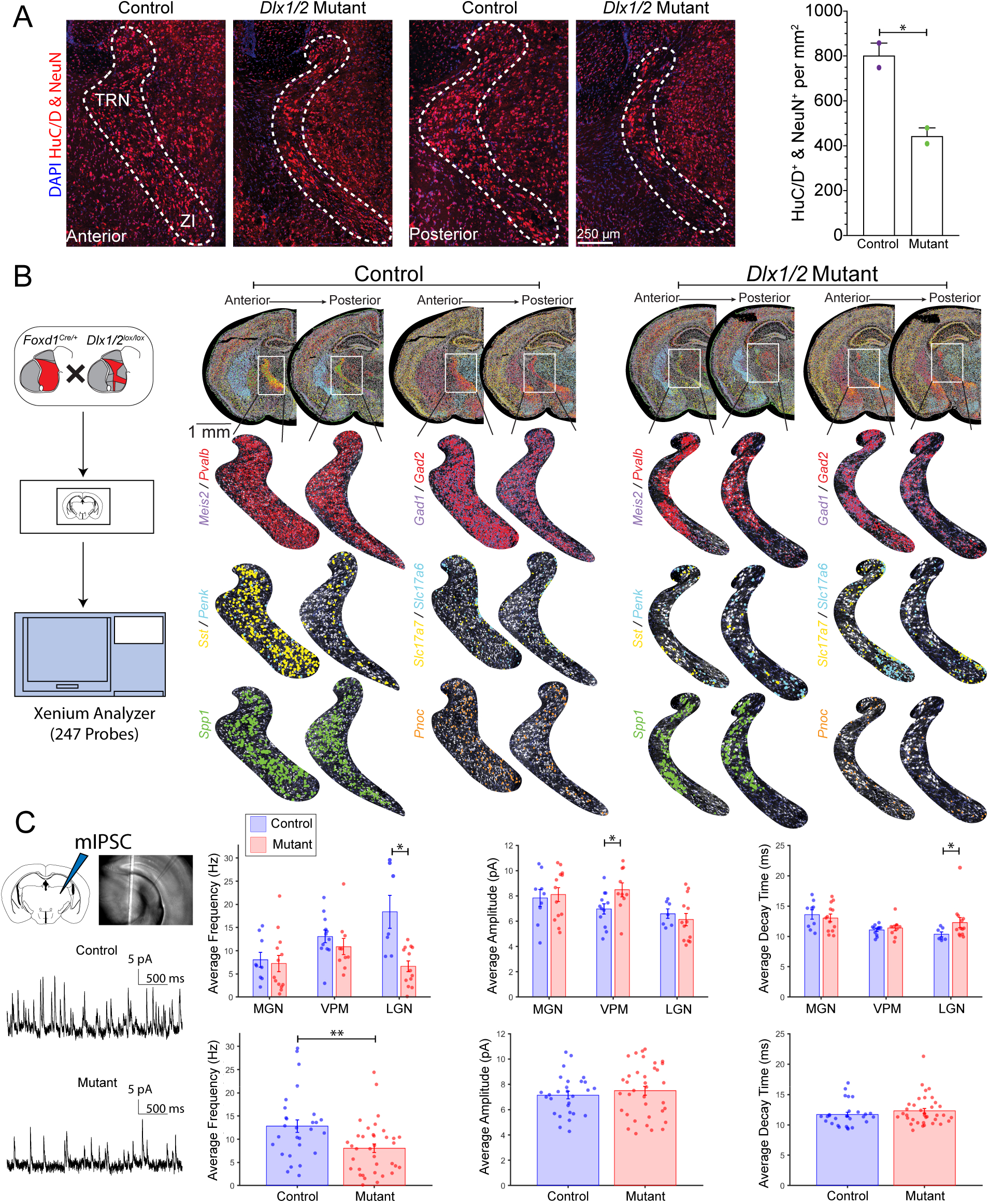
Dlx1/2 mediate transcriptional regulation of postnatal development and function of the thalamic reticular nucleus (TRN) and zona incerta (ZI) (additional data in Fig S12, S13, S14). **(A)** Reduced neuronal density in the TRN of *Dlx1/2* mutants. Left: immunostaining for DAPI, HuC/D, and NeuN (neuronal marker), in the anterior and posterior portion of the TRN and ZI. Dashed lines demarcate the boundaries of the TRN and ZI. Scale bar: 250 μm. Right: Quantification of HuC/D+ & NeuN+ neurons per mm² in the TRN. * *p < 0.05*, unpaired t-tests. **(B)** Spatial transcriptomics reveal disrupted gene expression in TRN and ZI: High-resolution Xenium spatial transcriptomics of controls and *Dlx1/2* mutants from anterior through posterior TRN regions. Differential expression is observed for markers such as *Meis2, Pvalb, Gad1, Gad2, Sst, Penk, Slc17a* and *Slc17a7*. Additional markers *Spp1* and *Pnoc* highlight changes in cellular composition across regions. **(C)** Miniature inhibitory postsynaptic current (mIPSC) recordings from the medial geniculate nucleus (MGN), ventral posteromedial nucleus (VPM), and lateral geniculate nucleus (LGN) of control and *Dlx1/2* mutant mice. Left: Schematic of recording site and representative mIPSC traces from control and mutant. Right: Quantification of mIPSC parameters: Average frequency, Average amplitude, and Average decay time across 3 brain regions (top row) or pooled analysis (bottom row) from control (blue) and mutant (red). Data are presented as mean ± SEM, with individual data points shown. \**p < 0.05*, \*\**p < 0.01*.

Consistent with predictions from GRN analysis (Fig S13A), *Sst*/*Penk*-expressing cells upregulated the adipogenesis-promoting factor *Arxes1*^65^(Tables S26, S27). Excitatory thalamic neurons, which are not derived from *Foxd1*-expressing diencephalic cells^66^, showed increased *Galanin* expression (Fig S12E, F, S13B, D). Additionally, glutamatergic neuronal markers *Slc17a6* and *Slc17a7* were elevated, particularly in the ZI (Fig 7B). Finally, a more than three-fold increase in non-neuronal cells was observed in the ZI and TRN (Fig S13E). This primarily reflects an increased abundance of Gfap-positive astrocytes, as no change in oligodendrocytes or microglia was observed (Fig S13F). Together, these findings suggest that *Dlx1/2* loss-of-function in the developing ZI and TRN disrupts the specification of GABAergic neurons.

### Loss of *Dlx1/2* in the Hypothalamus Leads to Hyperactivity, Sensory Deficits, and Impaired Inhibitory Inputs in the Thalamus

To investigate the consequences of *Dlx1/2* loss of function, we examined the behavioral phenotypes of gel-fed *Foxd1-Cre;Dlx1/2^lox/lox^*mutant mice compared to *Foxd1-Cre;Dlx1/2^lox/+^* controls. Mutant mice displayed hyperactivity, evidenced by significantly increased locomotor activity measured using DeepLabCut^67^ (Fig. S14A, Tables S30, S31). Mutant mice also spent less time in the periphery relative to the center of the open field, indicating reduced anxiety-like behavior (Fig. S14B, Table S32). Sensory function assays revealed heightened sensitivity to cold stimuli, as indicated by reduced response latency in the cold plate test, while responses to hot plate stimuli remained unchanged (Fig. S14C, Tables S33, S34). By contrast, motor coordination and learning assessed using the rotarod test showed no differences between mutants and controls (Fig. S14D, Tables S35).

These behavioral findings raised the hypothesis that *Dlx1/2* loss may impair ZI/TRN-mediated inhibitory circuits, resulting in impaired postsynaptic inhibition in the thalamus. To investigate the underlying neural basis of this behavior, we performed slice electrophysiology to assess spontaneous synaptic activity in the medial geniculate nucleus (MGN), ventral posteromedial nucleus (VPM), and lateral geniculate nucleus (LGN) of the thalamus (Fig 7C, S13G, Tables S36-S43). Whole-cell patch-clamp recordings showed that miniature excitatory postsynaptic currents (mEPSCs) did not differ in frequency, amplitude, or decay time between *Dlx1/2* mutants and control across all regions (Fig S13G). However, miniature inhibitory postsynaptic currents (mIPSCs) were markedly affected. Mutant mice exhibited a significant reduction in mIPSC frequency, particularly in the LGN (Fig 7C), accompanied by a modest increase in mIPSC decay time in the same region (Fig 7C). These deficits align with transcriptomic findings showing a substantial loss of inhibitory neurons, including parvalbumin-positive interneurons.

Together, these results indicate that *Dlx1/2* loss impairs inhibitory circuit function in the thalamus, potentially contributing to the observed hyperactivity and sensory processing deficits in *Dlx1/2* mutants.

These findings provide key insights into the regulatory roles of *Dlx1/2* in establishing and maintaining hypothalamic and prethalamic regional identity, as well as in guiding the differentiation of GABAergic versus glutamatergic neuronal populations. The observed disruptions in neurogenesis and neuronal composition in *Dlx1/2* mutants highlight the importance of these TFs in coordinating cell fate decisions that are critical for hypothalamic and prethalamic function and postnatal survival. More broadly, these results advance our understanding of how transcriptional regulatory networks control neuronal diversity and circuit formation within the hypothalamus and prethalamus, which may have implications for disorders involving dysfunctions in these brain regions.

## Discussion

In this study, we used integrated scRNA-Seq and scATAC-Seq analyses to map the GRNs governing the development of the hypothalamus and prethalamus in mice. This approach goes beyond previous scRNA-Seq analysis^24,26^ in providing a higher-resolution atlas of hypothalamic and prethalamic development, unveiling novel mechanisms that control neuronal subtype specification, and elucidating the dynamic interplay between transcriptional and chromatin states during neurogenesis. These insights contribute to a deeper understanding of how molecular mechanisms guide hypothalamic regionalization, neurogenesis, and the establishment of functional neural circuits. Integration of scRNA-Seq and scATAC-Seq across multiple developmental stages enabled the identification of temporally dynamic transcriptional states and active regulatory elements. The addition of scATAC-Seq-based chromatin accessibility profiling enabled us to delineate the transcriptional control mechanisms underpinning the establishment of distinct hypothalamic and prethalamic regions.

In addition to increasing the resolution of progenitor cell populations, we identified cross-repressive interactions among different classes of hypothalamic and prethalamic progenitors. This cross-repression particularly between prethalamic/tuberal progenitors and floorplate-like supramammillary/mammillary progenitors, limits progenitor plasticity and maintains regional boundaries. This is confirmed by analysis of function mutants of *Isl1*, *Nkx2-2*, and *Dlx1/2* – all of which are predicted to simultaneously promote prethalamic and/or tuberal identity while repressing supramammillary/mammillary identity. In each mutant line, prethalamic markers are lost while supramammillary/mammillary markers are upregulated. These findings extend earlier observations from the early stages of chick forebrain regionalization into later stages of hypothalamic and prethalamic neurogenesis^23^, and highlight conserved mechanisms regulating hypothalamic and prethalamic regional identity. The unexpected finding that genetic variants in transcription factors that promote supramammillary identity are associated a broad range of metabolic and behavioral traits highlight the importance of this relatively understudied hypothalamic region.

Our findings demonstrate that transcription factors that delineate regional identities and boundaries are not only structural but actively contribute to neuronal specification by regulating neuropeptide and neurotransmitter expression, therefore acting as ‘terminal selectors’^68,69^. For example, *Dlx1/2* were shown to be essential for specifying GABAergic and dopaminergic neuronal subtypes while also directly activating *Slc32a1* and *Th* expression. Importantly, many of these neuronal subtypes are specified before birth, suggesting that early gene regulatory programs have long-lasting effects on hypothalamic and prethalamic function. These early gene regulatory programs also provide a bridge between GWAS findings on neurodevelopmental disorders and both prethalamic and hypothalamic differentiation. The enrichment of multiple GWAS-implicated genes in neural progenitor cells of both the prethalamus and hypothalamus suggests that early disruptions in GRNs may underlie the pathology of neurodevelopmental disorders. For instance, genes linked to ASD, such as *Dlx1/2* and *Arx*, are critical for GABAergic neuron differentiation in the ZI and TRN^70–72^.

Analysis of the postnatal phenotypes of conditional *Dlx1/2* mutants highlights the importance of these factors in specifying GABAergic neurons in the ZI and TRN. Loss of *Dlx1/2* results in a reduction of the number of GABAergic neurons, and a corresponding reduction in postsynaptic inhibition in dorsal thalamic neurons. This alteration is associated with both hyperactivity and enhanced cold sensitivity in mutant mice, suggesting that disrupted development of inhibitory neurons in ZI/TRN may contribute to hyperactivity observed in ASD or ADHD^73,74^. Sensory hypersensitivity in ASD has likewise been linked to impaired inhibitory mechanisms in thalamic circuits^75^. ASD-related hyperactivity may also arise from thalamic dysfunction, resulting from an imbalance of excitatory-inhibitory regulation that disrupts normal motor and cognitive control^73,76,77^. The reduced GABAergic inhibition in *Dlx1/2* mutants parallels findings in other ASD models, where alterations in inhibitory signaling lead to increased thalamic and cortical excitability.

*Arx*, another ASD-related transcription factor^78–80^ which we identify as a direct target of Dlx1/2, also plays an essential role in prethalamic development^81,82^. This confirms that multiple TFs act in combination to regulate the excitatory-inhibitory balance in the ZI and TRN, which in turn affects thalamic function. Dysregulation of these networks likely contributes to neurodevelopmental disorders by impairing GABAergic neuron differentiation and disrupting thalamic excitatory-inhibitory homeostasis. This study highlights the potential relevance of defects in prethalamic patterning and neurogenesis to common neuropsychiatric disorders.

## Limitations of the Study

Although GRN analysis revealed robust regulatory relationships, several limitations should be noted. First, single-cell data, though powerful, may not fully capture gene interactions without experimental validation of knockout or knockdown approaches. Second, while this analysis ranked genes within GRNs, lower-ranked genes may still have functional contributions that were not fully explored. Third, repressive interactions were generally not as well-defined as their activating counterparts. Employing single cell CUT&TAG to identify targets of repressive factors may ultimately be essential for uncovering additional layers of regulatory complexity during hypothalamic development.

## Resource Availability

### Lead contract

Further information and requests for resources and reagents should be directed to and will be fulfilled by the Lead Contact, Seth Blackshaw.

### Materials availability

All unique/stable reagents generated in this study are available from the Lead Contact without restriction.

## Data and Code Availability

● All raw sequencing data, including scRNA-Seq and scATAC-Seq datasets, have been deposited in the NCBI Gene Expression Omnibus (GEO) repository (GSE284492).
● Processed Seurat objects and annotated datasets are hosted on Zenodo (10.5281/zenodo.14292831).
● Complete analysis code and workflows are available on GitHub (<u>github.com/dkim195/HyDD2</u>), ensuring reproducibility and transparency.
● Generated UMAPs will be available in the UCSC Cell Browser (Link pending)

## Supporting information

Reagents and Resources

## Acknowledgments

This work was supported by the NIH (R01MH126676) to S.B.; F31DK132944 from the National Institute of Diabetes and Digestive and Kidney Diseases (NIDDK) and F99-NS135816 from the National Institute of Neurological Disorders and Stroke (NINDS) to L.H.D.; RO1DC009607 from the National Institute of Deafness and Communication Disorders (NIDCD) to P.O.K.; and the Lundbeckfonden grant (R361-2020-2654), the Novo Nordisk Foundation grant (NNF24OC0089408), and the Parkinsonforeningen (R63-A1583-B890) to D.W.K.

We thank Transcriptomics and Deep Sequencing Core (Johns Hopkins) for sequencing of scRNA-Seq, scATAC-Seq, and Xenium libraries; Micfac Microscope Facility for providing infrastructure to perform imaging (Johns Hopkins); Behavioral Core for providing infrastructure to perform mouse behavior tests (Johns Hopkins); MARCC (Johns Hopkins) and GenomeDK (Aarhus University) for providing infrastructure to perform data analysis. *Isl1^lox/lox^* mice were a gift from Dr. Lin Gan, *Lhx1^lox/lox^* mice a gift from Dr. Richard R Behringer, and *Dlx1/2^lox/lox^* a gift from Dr. Jay Lee.

## Author Contributions

S.B., D.W.K., P.O.K., and E.P. conceived and supervised the study. D.W.K. and L.H.D. generated and analyzed sequencing data. J.X. and M.C. conducted and analyzed electrophysiology experiments. D.W.K. and L.H.D. performed and analyzed staining experiments. C.E.T., S.S.S., D.W.K., and L.H.D. conducted and analyzed behavioral experiments. D.W.K., L.H.D., and S.B. drafted the manuscript. All authors reviewed and edited the manuscript.

## Methods

### 1. Animal models

Time-mated wild-type (WT) CD1 mice were obtained from Charles River Laboratories. Transgenic mouse lines were maintained on a mixed C67BL/6 and CD1 background. The following transgenic mouse lines were used in the study:

1. *Foxd1-*Cre (B6;129S4-*Foxd1tm1(GFP/cre)Amc*/J, JAX#012463^83^),
2. *Isl1^lox/lox^* (gift from Dr. Lin Gan^84^),
3. *Lhx1^lox/lox^* (gift from Dr. Richard R Behringer^85^),
4. *Nkx2.2*-Cre (B6.129S6(Cg)-*Nkx2-2tm4.1(cre/EGFP)Suss*/J, JAX#026880^86^),
5. *Dlx1/2^lox/lox^* (gift from Dr. Jay Lee^52^).

Animals were housed in a temperature-controlled facility with a 14-hour dark/10-hour light cycle and had *ad libitum* access to food and water. All animal procedures followed the National Institutes of Health Guide for the Care and Use of Laboratory Animals and were approved by the Johns Hopkins University Animal Care and Use Committee.

### 2. Tissue Collection and Dissections

**Dissection of wildtype mice:** Tissues were collected from wildtype mice representing a random but equal mix of males and females across postnatal ages. For each embryonic stage analyzed, 10-30 embryos were harvested from 3-4 pregnant dams, depending on the gestational age. Brain dissections followed previously established protocols^24^. Meninges were removed, and dissections were completed in cold HBSS (#14170112, ThermoFisher Scientific) within 30 minutes for each age group.

**Dissection of mutant mice:** Embryos from mutant mice were collected from a minimum of five pregnant dams. Hypothalamic and prethalamic regions were dissected, and the remaining tissues of embryos were used for rapid genotyping (#4359187, ThermoFisher Scientific), and grouped by genotype for further analysis. Dissected hypothalamic and prethalamic tissues were stored in HABG buffer (Hibernate-E (#A1247601, ThermoFisher Scientific) for embryos or Hibernate-A (#A1247501, ThermoFisher Scientific) for postnatal stages, both supplemented with 2% B-27 Supplement (#17504044, ThermoFisher Scientific), and 0.5 mM GlutaMAX Supplement (#35050061, ThermoFisher Scientific) until cell dissociation.

### 3. Preparations of Cells and Nuclei

**Cell dissociation for scRNA-Seq:** To prepare samples for scRNA-Seq, tissues were dissociated in Hibernate medium (Hibernate-E Minus Calcium (#HECA500, BrainBits) for embryos or Hibernate-A Minus Calcium (#HACA500, BrainBits) for postnatal stages), supplemented with 2 mg/ml papain (#LS003119, Worthington Biochemical) and 0.5 mM Glutamax. The enzyme mixture was pre-incubated for 15 minutes, then added to tissues for 5 - 30 minutes with 0.1 U/μl RNase inhibitor (N2615, Promega). Toward the end of digestion, 100 μg/ml final DNase I (#4716728001, Sigma-Aldrich) was added. Cells were subsequently centrifuged, resuspended in HABG buffer, filtered through a 40 μm cell strainer (BAH136800040-50EA, Sigma-Aldrich), and resuspended in HABG buffer containing RNase inhibitor (0.5 U/μl). Cell viability and counts were assessed with Trypan Blue staining (#15250061, Thermo Fisher Scientific) and a hemocytometer.

**Nuclei preparation for scATAC-Seq:** Nuclei for scATAC-Seq were extracted following post-cell dissociation and purified following a 10x Genomics protocol. Nuclei count and yield were verified with Trypan Blue staining and a hemocytometer.

**Nuclei preparation for snRNA-Seq:** For snRNA-Seq, adult mouse brains were rapidly dissected, and 100 μm coronal sections were prepared using a mouse brain matrix. Regions spanning the ZI, TRN, and thalamus were microdissected under a dissection microscope. Nuclei were extracted following a 10x Genomics protocol.

### 4. Sequencing Library Preparation and Data Processing

**Sc/snRNA-Seq:** Libraries for scRNA-Seq and snRNA-Seq were prepared using the 10x Genomics Chromium. Single Cell 3’ v3.0 and v3.1 platforms (Table S1), following the manufacturer’s protocols. Sequencing was conducted on an Illumina NovaSeq 6000, targeting a depth of 500 million reads per sample. Raw data were processed with the 10x Genomics Cell Ranger Software (v5) and aligned to the mm10 mouse genome reference (refdata-gex-mm10-2020-A).

**scATAC-Seq:** scATAC-Seq libraries were prepared using the 10x Genomics Chromium Single Cell ATAC v1.1 platform (Table S1), with one-hour tagmentation and 9-10 cycles of PCR. Libraries were sequenced on Illumina NovaSeq 6000, at a target depth of 600 million reads per sample. Raw data were processed with Cell Ranger ATAC (v2) and aligned to the mm10 mouse genome reference (refdata-cellranger-arc-mm10-2020-A-2.0.0).

1. 5. *In Situ* Hybridization

**DIG *in situ* hybridization:** DIG-labeled probes were used for *Dlx2*, *Isl1*, *Arx*, *Foxb1*, *Dlx5*, *Foxa1*, *Gad1*, and *Hmx2* mRNA detection, with colorimetric detection using NBT/BCIP substrates, using standard experimental protocols^24,30^.

**Single-molecule fluorescent *in situ* (smfISH) hybridization:** Mouse embryos were immersion-fixed and adult mouse brains were perfusion-fixed with 4% paraformaldehyde (PFA). Adult mouse brains were post-fixed in PFA for an additional 24 hours at 4°C. After fixation, tissues were cryoprotected in 30% sucrose solution, and embedded in an Optimal Cutting Temperature (OCT) compound. Brains were sectioned coronally at a thickness of 20 μm using a cryostat and mounted onto Superfrost Plus glass slides (#12-550-15, ThermoFisher Scientific). The tissue sections were stored at −80°C until further processing.

RNAscope (Advanced Cell Diagnostics, ACD) was used for smfISH, with the RNAscope^TM^ V2 Assay standard protocol as provided by the manufacturer. The following target-specific RNAscope probes were used: *Sp8* (Mm-Sp8-C3, #547521-C3), *Sp9* (Mm-Sp9-C1, #564531), *Pitx2* (Mm-Pitx2-C2, #412841-C2), *Irx5* (Mm-Irx5-C1, #513871), *Meis2* (Mm-Meis2-C2, #512841-C2), *Npw* (Mm-Npw-C1, #495671), *Pnoc* (Mm-Pnoc-C2, #437881-C2), *Slc32a1* (Mm-Slc32a1-C2, #319191-C2), *Slc17a6* (Mm-Slc17a6-C3, #319171-C3), and *Gal* (Mm-Gal-C1, #400961).

Sections were stained with DAPI, coverslipped using ProLong Gold Antifade Mountant (#P36934, Thermo Fisher Scientific), and stored at 4°C in the dark until imaging. Fluorescent images were acquired using a Zeiss LSM 780 confocal microscope. Multiple regions of interest were imaged per section, capturing different regions in embryos and adult brains.

### 6. Immunostaining

Brains were prepared as described above. Immunostaining was performed following the previous protocol^87^, using primary antibodies for Pvalb (Rabbit polyclonal, Swant PV-25, 1:400) and Pax6 (Rabbit polyclonal, ab2237, 1:200). Sections were incubated with primary antibodies overnight at 4°C, followed by a one-hour secondary antibody incubation.

### 7. Xenium *In Situ* Spatial Transcriptomics

P21 *Foxd1^Cre/+^;Dlx1/2^lox/+^* and *Foxd1^Cre/+^;Dlx1/2^lox/lox^* mice were anesthetized with 1 mL of tribromoethanol/Avertin and perfused transcardially with 1xPBS to clear blood from brain tissue. Fresh Frozen samples were prepared by flash-freezing dissected brains in an OCT embedding compound, following the 10x Genomics Xenium Fresh Frozen Tissue preparation protocol (CG000579, 10x Genomics). Four 12 μm coronal brain sections, spaced 300 μm apart, were taken from regions spanning from the preoptic area to the mammillary region and positioned on Xenium slides. Hybridization, ligation, and amplification were performed according to the 10x Genomics Xenium *In Situ* protocol (CG000582) with a panel of 100 custom gene probes and a mouse brain gene panel containing 247 probes (Table S28).

**Post-Xenium Immunohistochemistry:** Following Xenium imaging, slides were washed twice in 1xPBS, and incubated in a blocking buffer (0.4% Triton X-100, 10% Horse Serum in 1x PBS) for 2 hours at room temperature. Primary antibodies for HuC/D (mouse monoclonal, A-21271, 1:100), and NeuN (mouse monoclonal, MAB377, 1:100) in blocking buffer were added, and slides were incubated overnight at 4°C. Following primary antibody incubation, slides were washed twice in 1xPBS and incubated with a Goat anti-Mouse Alexa Fluor 647 (ab150115, 1:1000) in a blocking buffer for 1 hour at room temperature. Slides were then washed twice in 1x PBS and incubated with DAPI (1:1000) in 1xPBS for 5 minutes at room temperature. Sections were mounted using Aqua-Poly/Mount (#18606-20, Fisher Scientific) and coverslipped. Imaging was conducted on a Zeiss Axio Observer with an Apotome III Microscope (20x objective lens, 0.8 NA). Images were analyzed using QuPath and Xenium Explorer (v3).

### 8. Mouse Monitoring

**Video Monitoring in the Mouse Home-cage:** *Foxd1^Cre/+^;Dlx1/2^lox/+^*and *Foxd1^Cre/+^;Dlx1/2^lox/lox^* mice activities were monitored with a high-definition video camera positioned above the cage, capturing detailed movements. The camera was set to record at 60 frames per second in a quiet environment with a consistent lighting environment. Each mouse was recorded for one hour, and the footage was analyzed using DeepLabCut for behavior quantification.

**Survival Curve Monitoring and Analysis:** Two experimental groups, *Foxd1^Cre/+^;Dlx1/2^lox/+^* and *Foxd1^Cre/+^;Dlx1/2^lox/lox^* mice, were housed in groups of 3-5 animals per cage, with *ad libitum* access to standard food pellets and water. Additionally, an experimental subset of *Foxd1^Cre/+^;Dlx1/2^lox/lox^*mice received only liquid gel food and hydration diets, as this genotype exhibited significantly reduced size compared to the *Foxd1^Cre/+^;Dlx1/2^lox/+^*mice. Survival was monitored twice weekly starting from P21. Survival data, including survival rates, dates of death, and other relevant parameters, were recorded to evaluate the effects of genotype and dietary conditions on survival outcomes.

### 9. Behavior Assays

Behavior assays were conducted between 10 am and 2 pm over four days, using both male and female *Foxd1^Cre/+^;Dlx1/2^lox/+^*and *Foxd1^Cre/+^;Dlx1/2^lox/lox^* mice of the P21-P25 for each test. Day 1: Hot/Cold Plate assay was conducted, and the Rotarod test was initiated. In the Rotarod test, each mouse was given three trials per day with a 2-minute inter-trial interval, continuing over three consecutive days. Day 4: An Open Field Test was conducted.

Data analysis and graphing were performed using GraphPad Prism 10. The Open Field Test and Hot/cold Plate test results were analyzed with an unpaired T-Test, while one-way ANOVA with multiple comparisons was used for Rotarod Data.

**Open Field Test:** To assess locomotor activity and anxiety-like behavior, mice were placed in a 40 x 40 cm activity chamber equipped with infrared beams (San Diego Instruments Inc.). Total activity and time spent in the center of the chamber were automatically recorded with ‘Photobeam Activity System - Open Field (San Diego Instruments Inc.) over 30 minutes.

**Rotarod Test:** Motor coordination and learning were assessed using an accelerating rotarod (Columbus Instruments), starting at 4 RPM and accelerating at 7.2 RPM/minute. The latency and speed at which each mouse fell were recorded for each trial.

**Hot/Cold Plate Test:** Thermal sensitivity was measured using the hot/cold plate test. Mice were placed on a 16.5 x 16.5 cm metal plate (Bioseb, FL, USA) heated to 54°C for the hot plate test or cooled to 4°C for the cold plate test. Latency to hind limb withdrawal was recorded, with a 30-second cut-off if no response occurred. Mice were immediately returned to their home cage following the test.

### 10. Slice Electrophysiology

**Preparation of brain slices:** *Foxd1^Cre/+^;Dlx1/2^lox/+^*and *Foxd1^Cre/+^;Dlx1/2^lox/lox^* mice were deeply anesthetized with isoflurane prior to decapitation. The brain was quickly removed and submerged in an ice-cold dissection solution containing (in mM): 212.7 sucrose, 2.6 KCl, 1.23 NaH_2_PO_4_, 26 NaHCO_3_, 10 glucose, 3 MgCl_2_.6H_2_O, and 1 CaCl_2_.2H_2_O (pH 7.35–7.4, equilibrated with 95% O_2_–5% CO_2_). The brain was blocked and sliced coronally using a slicing microtome (Leica VT1200) in the ice-cold dissection solution. Coronal slices (400 μm thickness) containing either the medial geniculate nucleus (MGN), lateral geniculate nucleus (LGN), or ventral posteromedial nucleus (VPM) were obtained. Slices were then incubated in oxygenated artificial cerebrospinal fluid (ACSF, in mM): 130 NaCl, 3 KCl, 1.25 NaH_2_PO_4_, 20 NaHCO_3_, 10 glucose, 1.3 MgSO_4_.7H_2_O, and 2.5 CaCl_2_.2H_2_O (pH 7.35–7.4, equilibrated with 95% O_2_–5% CO_2_) for 50 minutes at 30°C. After incubation, the slices were maintained at room temperature until recording.

**Electrophysiological recording:** Slices were placed in a recording chamber on a fixed-stage microscope (Olympus BX51) and superfused with oxygenated ACSF at 2-4 mL/min at room temperature. Tetrodotoxin (TTX, 1 μM) was added to the recording solution to block the generation and propagation of action potentials. Whole-cell recordings were performed using recording pipettes with an input resistance of 4-12 MΩ filled with a cesium-based internal solution containing (in mM): 115 cesium methanesulfonate (CsCH_3_SO_3_), 5 NaF, 10 EGTA, 10 HEPES, 15 CsCl, 3.5 MgATP, and 3 QX-314 (pH 7.25; 300 mOsm). The membrane potential was clamped at - 70 mV and 0 mV to record miniature excitatory and inhibitory postsynaptic currents (mEPSCs and mIPSCs), respectively. After a stable whole-cell recording was established, recordings were performed for 10 minutes at each holding potential. Voltages were corrected for an estimated junction potential of 10 mV. All recordings were performed using a patch clamp amplifier (MultiClamp 700B, Molecular Devices), digitized using a DAQ board (NI PCI-6259, National Instruments), and acquired using the Wavesurfer software (HHMI Janelia Farms, https://wavesurfer.janelia.org).

### 11. Data Analysis

All codes used in this work are publicly available at <u>github.com/dkim195/HyDD2</u> and sample details are available in Table S1. Below, we briefly describe the methodologies employed for the analyses presented in this study.

### 11A.#Single-cell Sequencing

**ScRNA-Seq:** scRNA-Seq data were processed using Seurat v3.2.3. Cells with fewer than 1,000 detected genes, fewer than 2,000 unique molecular identifiers (UMIs), or more than 50% mitochondrial transcript content were excluded. Additionally, cells with over 25% ribosomal transcript content were removed. To minimize the impact of technical artifacts, doublets were identified and excluded based on prior analysis of matched datasets^24^.

The dataset was normalized using SCTransform, regressing out the effects of the number of genes (*nFeature_RNA*) and UMIs (*nCount_RNA*). Dimensionality reduction was performed using principal component analysis (PCA). To visualize clustering results, uniform manifold approximation and projection (UMAP) embeddings were computed using the top 20–30 principal components. Clustering was performed using the Louvain algorithm implemented in Seurat’s *FindClusters()* function, with the resolution parameter optimized for biological relevance.

Clusters were annotated through an iterative process. In the first pass, clusters were assigned based on cell-type-specific markers, and contaminant populations such as immune cells, endothelial cells, and meningeal cells were removed. Subsequently, datasets from each developmental stage were merged, and clusters were rechecked and refined to ensure consistency. After integrating data from multiple developmental stages, ambiguous clusters such as those containing mixed progenitor and immune cell populations were labeled as “check” clusters for further examination. Batch effects arising from technical and biological sources were addressed using Harmony^88^, which corrected for variability associated with sample origin (*orig.ident*) and sex-linked gene expression (e.g., *Xist* and *Ddx3x*). This integrated dataset allowed seamless comparisons across developmental stages and genotypes.

Cluster-specific marker genes were identified using the Wilcoxon rank-sum test implemented in the *FindMarkers()* function. Differential gene expression (DEG) analysis was performed with a minimum log2 fold change threshold of 0.5 and a minimum fraction of 20% of cells expressing the gene. Enrichment analysis of the resulting DEG lists was used to identify transcription factors driving lineage-specific programs.

**ScATAC-Seq:** scATAC-Seq data underwent similar quality control measures to scRNA-Seq data. In general, nuclei with fewer than 1,000 detected peaks or 2,000 fragments were excluded from the analysis (Table S1). Chromatin accessibility data were normalized using SCTransform, and dimensionality reduction was performed using PCA. UMAP embeddings were computed after correcting for batch effects using Harmony, with corrections applied to account for sample origin (*orig.ident*).

Clusters were identified using the Louvain algorithm and annotated through an iterative refinement process. Initial low-quality clusters, such as those with high mitochondrial content, were excluded. Following this, data from multiple developmental stages were merged, and cluster annotations were refined to ensure consistency across samples. To better understand lineage-specific chromatin accessibility, differential accessibility analysis was performed using the *FindMarkers()* function, with a log2 fold change threshold of 0.5 and a requirement that peaks be accessible in at least 20% of cells. Motif enrichment analysis was used to identify transcription factors associated with differentially accessible regions.

**Integration of scRNA-Seq and scATAC-Seq Data:** Integration of scRNA-Seq and scATAC-Seq datasets was performed to annotate scATAC-Seq cell types based on transcriptional profiles. To achieve this, cell-type labels from scRNA-Seq data were transferred to matching scATAC-Seq datasets using Seurat’s *Transfer Anchor* workflow. Annotation was guided by known cell-type-specific marker genes and chromatin accessibility profiles at their respective loci. This approach enabled the alignment of transcriptomic and epigenomic data, facilitating a comprehensive analysis of cell-type-specific gene regulation.

**Developmental Progression and Mutant Atlas:** For developmental progression analyses, scRNA-Seq and scATAC-Seq datasets were used to identify changes in cell-type composition, transcriptional programs, and chromatin accessibility across different developmental stages. Specific attention was paid to early patterning, neurogenesis, gliogenesis, and neural specification.

To explore mutant phenotypes, an atlas was generated using scRNA-Seq and scATAC-Seq datasets from E12.5 wild-type and mutant samples. scATAC-Seq data were aligned to their scRNA-Seq counterparts, enabling the transfer of cell-type annotations. To ensure consistency, scATAC-Seq datasets were downsampled to match the number and quality of cells in the scRNA-Seq datasets. Differential analysis between wild-type and mutant datasets was performed to identify mutant-specific changes in gene expression and chromatin accessibility.

### 11B.#SCENIC+ Gene Regulatory Network Analysis

**Preparation of scRNA-Seq Data for SCENIC+:** Single-cell RNA sequencing (scRNA-Seq) data were processed and converted for SCENIC+ analysis^27^ using Seurat (v4.3.0)^89^ and Scanpy (v1.9.1)^90^. For each dataset, cell clusters annotated through earlier steps were retained, while cells identified as doublets or those from non-relevant clusters (e.g., high mitochondrial content or non-neuronal clusters) were excluded. The Seurat objects for different datasets—patterning, neurogenesis, and mutant samples were converted to AnnData format using the *MuDataSeurat::WriteH5AD* function.

Metadata, including sample origin, counts, features, cluster identity (Cluster_Pass2), and genotype, were preserved. UMAP was exported as CSV files and integrated into AnnData objects using Scanpy. Raw counts were stored in a separate layer to retain unnormalized data for downstream SCENIC+ processing. Gene expression data were normalized to a total count of 10,000 and log-transformed prior to SCENIC+ processing. The prepared AnnData objects were saved in the h5ad format for subsequent integration with scATAC-Seq data.

**Preparation of scATAC-Seq Data for SCENIC+:** scATAC-Seq data were processed using Signac (v1.9.0) and PycisTopic. Initial quality control steps included the removal of low-quality cells, cells identified as doublets, and those originating from non-brain regions. Clusters annotated as “NPC (Glia)” were downsampled to 10% of their original size to balance representation in the dataset.

Peak count matrices were converted to the Feather format using the *arrow* package. Metadata, including cluster annotations (Cluster_Pass2) and UMAP embeddings, were exported alongside the count matrices. A PycisTopic object was created, integrating count matrices, cell annotations, and fragment file paths for each sample. MALLET-based topic modeling was performed using 50 topics with 500 iterations, employing alpha and eta hyperparameters for Dirichlet distributions. The resulting topic models were evaluated to identify the most optimal model based on log-likelihood scores. UMAP embeddings were generated from the PycisTopic object and aligned with RNA-based UMAP embeddings.

**Integration of scRNA-Seq and scATAC-Seq Data in SCENIC+:** For SCENIC+ analysis, scRNA and scATAC datasets were integrated using the SCENIC+ Snakemake pipeline. AnnData objects containing RNA expression data were paired with cisTopic objects containing scATAC-Seq accessibility data. Regions-to-genes associations were inferred using proximity rules and co-accessibility relationships. Direct and extended regulatory regions were defined based on motif annotations and overlap with chromatin accessibility peaks.

For motif enrichment, we used the v10nr_clust motif collection, with annotations specific to *Mus musculus*. Differential motif enrichment analysis was conducted using adjusted p-values < 0.05 and log2FC thresholds ≥ 1.0 to identify transcription factors (TFs) enriched within cell-type-specific accessible regions. To evaluate TF-to-gene relationships, GEX data were modeled using the Gradient Boosting Machine algorithm. Regulatory modules (eRegulons) were identified and scored using AUCell, with both direct and extended annotations.

To identify differentially accessible regions, pseudobulk peak calling was performed for each cluster. Barcode-to-fragment mappings were generated for each cell, and pseudobulk peaks were called using MACS2 with parameters optimized for scATAC-Seq data (e.g., shift = 73 bp, extension = 146 bp). Consensus peaks were determined by merging narrow peaks across samples and filtering against the mm10 blacklist. The resulting peaks were annotated using Ensembl genome annotations.

The SCENIC+ pipeline was implemented using a Snakemake workflow. Configurations included genome annotation, chromsizes, and motif annotation files. The integrated SCENIC+ pipeline generated eRegulons for each cluster, which were used to infer cell-type-specific regulatory landscapes and to identify key transcriptional regulators of neurogenesis and patterning. Final outputs included inferred regulatory networks, motif enrichment results, and direct/extended eRegulons, which were stored in H5Mu and TSV formats for downstream interpretation.

**Identification and Annotation of eRegulons:** GRNs were constructed using the SCENIC+ pipeline by integrating single-cell RNA-Seq and ATAC-Seq data. To identify eRegulons, the extended regulatory regions were analyzed for transcriptional activation and repression motifs. The SCENIC+ pipeline inferred TF-to-gene regulatory interactions through motif enrichment analyses, evaluating the importance of TF-to-gene relationships. Extended eRegulons were filtered for significant activator (extended +/+) or repressor (extended -/+) motifs, and genes with high TF-to-gene importance scores and ranks were retained for downstream analyses.

Gene sets representing distinct hypothalamic domains and developmental stages were curated based on prior knowledge. These included genes for the supramammillary nucleus (SMN), medial mammillary nucleus (MMN), paraventricular and supraoptic nuclei (PVH_SON), tuberal regions (Tub), premammillary nucleus (PMN), and prethalamus (PreThal), as well as progenitor-specific markers. Genes were grouped into activator or repressor types, based on motif enrichment within eRegulons, and cross-referenced with experimentally validated transcription factors (TFs) obtained from public databases. Only TFs with significant eRegulon associations and non-conflicting annotations were included in further analysis.

To identify feedback regulatory interactions, custom R functions were implemented to detect pairs of TFs and target genes that formed reciprocal or overlapping regulatory relationships within eRegulons. Positive feedback loops were defined as activator-activator interactions, while negative feedback loops consisted of activator-repressor or repressor-repressor pairs. Each regulatory relationship was labeled with its origin cluster and type of feedback (activator or repressor). Feedback regulatory networks were constructed for specific clusters, such as SMN, MMN, and progenitor domains, and later integrated into a combined network across clusters. The filtered GRN included only high-confidence TF-target relationships, which were grouped by hypothalamic domains or cell types.

Gene regulatory networks were visualized using the igraph and ggraph R packages, which were refined further using Cytoscape (v3.10.3). Regulatory relationships were plotted as directed graphs, where nodes represented TFs or target genes and edges denoted activation or repression interactions. Feedback loops and cross-cluster regulatory connections were highlighted. Edge thickness was scaled by the TF-to-gene importance score, while edge color represented activator (blue) or repressor (red) interactions. Nodes were annotated with their respective clusters or domains, and network layouts were optimized using force-directed algorithms.

To further analyze interactions between hypothalamic regions, regulatory relationships were aggregated by their source (TF origin cluster) and target (regulated cluster). Region-level interaction networks were visualized with directional edges representing activation or repression between clusters. Specific subnetworks were extracted to examine interconnectivity between progenitor domains and terminally differentiated cell types, such as SMN and MMN.

### 11C.#Xenium Spatial Transcriptomics Analysis

**Data Acquisition and Preprocessing:** Raw Xenium data were imported into the *SpatialData* framework via the *spatialdata_io module*, ensuring compatibility with downstream analyses. Raw spatial datasets were converted to *.zarr* format and processed further using *Scanpy* and *Squidpy*.

Cellular quality metrics, including total transcripts, unique transcripts, segmented cell areas, and nucleus-to-cell area ratios, were calculated for each region using *scanpy.pp.calculate_qc_metrics*. Cells with low transcript counts or other aberrant quality metrics were flagged for potential exclusion. Negative control DNA probe counts and decoding counts were assessed to ensure data integrity.

**Data Normalization and Dimension Reduction:** For each region, raw transcript count matrices were normalized using total count normalization (*scanpy.pp.normalize_total*) and subsequently log-transformed (*scanpy.pp.log1p*). Highly variable genes were identified (*scanpy.pp.highly_variable_genes*) for downstream clustering and differential expression analysis. PCA was performed to reduce dimensionality, retaining components that captured the majority of variance.

**Clustering and Neighborhood Analysis:** Spatially resolved clusters were identified using the Leiden clustering algorithm (*scanpy.tl.leiden*) applied to PCA-derived neighbors. To assess spatial relationships among clusters, spatial neighbors and neighborhood enrichment were calculated (*squidpy.gr.spatial_neighbors* and *squidpy.gr.nhood_enrichment*, respectively). Enrichment matrices were visualized to explore cluster co-occurrence patterns.

**Visualization of Spatial Gene Expression:** Clusters and individual gene expression levels were visualized in a spatial context using *squidpy.pl.spatial_scatter*.

### 11D.#Behavioral Analysis Using DeepLabCut

A DeepLabCut (DLC)^67^ project was created for the automated tracking of movement in behavioral videos. The project, titled *“Dlx“*, was initialized using the *deeplabcut.create_new_project* function, specifying the experimenter, video file paths, and project settings. The system was configured for single-animal tracking with anatomical landmarks including Head, Neck, and Tail, as determined via the graphical user interface. Additional videos were imported as required during the project lifecycle.

Key frames were automatically extracted from the videos using the k-means clustering algorithm via *deeplabcut.extract_frames* (mode = “automatic“). Extracted frames were manually labeled for the specified landmarks to ensure accurate identification.

A training dataset was generated using the labeled frames and augmented automatically by DLC to improve model robustness. This step included splitting datasets into training and validation subsets via the GUI.

The neural network was trained using the labeled data. Model training included the optimization of parameters to achieve accurate landmark detection. Post-training, the network’s performance was evaluated with validation datasets using *deeplabcut.evaluate_network*. Performance metrics and visualized results were generated, providing insight into the model’s accuracy.

Trained models were deployed to analyze new videos using *deeplabcut.analyze_videos*. Key options such as saving results as CSV files, filtering predictions, and visualizing trajectories were configured to enhance tracking reliability. For analysis of specific videos, paths and settings were programmatically defined, enabling automated processing.

To improve tracking precision, trajectory data were post-processed using DLC’s tracking and refinement tools. The detected movement was converted into tracklets with the *deeplabcut.convert_detections2tracklets* function. Tracklets were further stitched (*deeplabcut.stitch_tracklets*) to produce continuous trajectories, filtered (*deeplabcut.filterpredictions*) for noise reduction, and visualized (*deeplabcut.plot_trajectories*).

Labeled videos with trajectories were generated for quality control and presentation using *deeplabcut.create_labeled_video*. Visualizations included color-coded markers, skeleton overlays, and trailing trajectories, highlighting tracked movements.

When necessary, outlier frames were extracted and manually refined via the GUI. Updated labels were merged back into the dataset (*deeplabcut.merge_datasets*), followed by retraining the model (*deeplabcut.train_network*) with a new shuffle index to incorporate refined annotations.

Analysis of the output was conducted following github.com/ETHZ-INS/DLCAnalyzer.

### 11E.#Electrophysiological Recording Analysis

Electrophysiological data were analyzed using the MATLAB-based data analysis software ‘minis’^91^. Recording traces were initially band-pass filtered at 50 Hz and smoothed using a 1.5-millisecond Gaussian window to remove high-frequency noise. Events were identified in the smoothed trace, and each event was evaluated using an amplitude threshold of 3 pA and a maximum time-to-peak of 5 milliseconds. All data were plotted with means ± standard error of the mean (SEM). Data normality was assessed using the Shapiro-Wilk (SW) test and comparisons were made using either a Student t-test or rank-sum test, as appropriate.

### 12. Statistics

Statistical analyses were conducted using GraphPad Prism or R (codes available in GitHub). Data are presented as the mean ± SEM. A P-value of less than *0.05* was considered statistically significant. Specific details of each statistical test are provided in the relevant method section.

## Supplemental Figures

Figure S1. Transcriptional and chromatin accessibility dynamics during hypothalamic development (related to Figure 1).

**(A)** *Patterning & Neurogenesis* phase. Top: UMAP of scRNA-Seq data from E11–E14 hypothalamic cells, displaying distinct clusters of neural progenitor and early neuronal populations. Bottom: UMAP of scATAC-Seq data, showing motif accessibility for key transcription factors associated with early hypothalamic patterning.

**(B)** *Differentiation* phase. Top: UMAP of scRNA-Seq data from E15–P8 hypothalamic cells, highlighting the emergence of mature neuronal populations, including tanycytes, astrocytes, oligodendrocytes, and specific hypothalamic nuclei. Bottom: Feature plots showing expression of key hypothalamic genes, such as *Agrp*, *Npvf*, *Tac1*, and *Slc30a3*. Inset displays *in situ* hybridization from the Allen Brain Atlas of *Slc30a3* in the Zona Incerta.

**Figure S2. Gene expression analysis during hypothalamic patterning and neurogenesis (related to Figure 1).**

**(A)** UMAP projection of scRNA-Seq data from E11 to E14 hypothalamic cells, showing distinct clusters corresponding to neural progenitor and differentiating neuronal populations.

**(B)** Top: UMAP plot showing expression of *Bsx*, a transcription factor. Bottom: *in situ* hybridization from the Allen Brain Atlas at E13.5 confirms *Bsx* expression in the putative suprachiasmatic nucleus (SON) (dashed line); sagittal section. Scale bar: 200 μm.

**(C)** Feature plots showing expression patterns of select transcription factors and neuropeptides across hypothalamic progenitors and neuronal subtypes.

**Figure S3. Gene regulatory networks and transcription factor dynamics during hypothalamic development (related to Figure 2).**

**(A)** Schematic of SCENIC+ analysis applied to hypothalamic scRNA-Seq and scATAC-Seq data from the *Patterning & Neurogenesis* phase (E11–E14). This approach identifies gene regulatory networks (GRNs) composed of transcription factors (TFs) that function as activators (blue arrows) or repressors (red arrows).

**(B)** UMAP plot showing the distribution of hypothalamic progenitor populations.

**(C)** Heatmap displaying relative expression (z-scores) of key TFs across hypothalamic progenitors and early neuronal populations.

**Figure S4. Integration of hypothalamic gene regulatory networks with EMBI-EBI GWAS traits (related to Figure 2).**

**(A)** Schematic of SCENIC+ analysis applied to hypothalamic scRNA-Seq and scATAC-Seq data, integrated with GWAS traits from the EMBI-EBI database.

**(B)** Heatmap displaying correlations between hypothalamic regionally restricted TFs and various disease traits, with z-scores indicating the strength of association.

**Figure S5. Gene regulatory networks and transcription factor dynamics during hypothalamic cell type differentiation (related to Figure 3).**

**(A)** Overview of SCENIC+ analysis applied to scRNA-Seq and scATAC-Seq data from E15 to P8, identifying transcription factors (TFs) driving the specification of distinct hypothalamic cell types. GRNs show TFs acting as activators (blue arrows) or repressors (red arrows).

**(B)** UMAP plot showing hypothalamic cell populations.

**(C)** Heatmap displaying relative expression (z-scores) of key TFs across hypothalamic cell types.

**Figure S6. Identification of TH-expressing clusters and their regulatory networks in the hypothalamus.**

**(A)** UMAP projections showing *Th*-expressing clusters within the hypothalamus, including prethalamic clusters (PreThal1, PreThal2, PreThal3), paraventricular hypothalamic nucleus (PVH/SON1, PVH/SON2), and distinct subpopulations in the premammillary nucleus (PMN_Nts, PMN_Gal, PMN_Ghrh), along with other regions such as the supramammillary nucleus (SMN) and ISL1+ clusters.

**(B)** Gene regulatory network (GRN) identifying key transcription factors predicted to drive *Th* expression, including *Isl1*, *Dlx*, *Meis*, and *Hmx* family members.

**(C)** Feature plots showing the expression of additional markers in *Th*-expressing clusters, including *Arx*, *Esr1*, *Trh*, *Tac1*, *Gal*, *Prlr*, *Npw*, *Npy*, *Foxa2*, *Ghrh*, *Gpr101*, and *Nts*.

**Figure S7. Disruption of *Lhx1* affects gene regulatory networks and neuronal identity in the hypothalamus (related to Figure 4).**

**(A)** Violin plot showing reduced *Lhx1* expression in *Lhx1* mutants compared to controls at E12.5.

**(B)** UMAP projections comparing the distribution of progenitor populations in control and *Lhx1*-mutant hypothalamus.

**(C)** Feature plots showing the expression of key neuronal markers, including *Ascl1*, *Neurog2, Foxb1, Nr4a2, Sim1, Shox2, Foxa2, Calb2, Lmx1b,* and *Barhl2*, in control and *Lhx1*-mutant hypothalamus.

**Figure S8. Impact of *Isl1* disruption on gene regulatory networks and hypothalamic progenitors (related to Figure 4).**

**(A)** Top: violin plot showing reduced *Isl1* expression in *Isl1* mutants compared to controls at E12.5. Bottom: expression of *Foxa1* and *Foxb1* in control versus *Isl1* mutant hypothalamus.

**(B)** Top: UMAP projections from scATAC-Seq data comparing control and *Isl1* mutants, showing changes in the clustering of progenitor populations. Bottom: chromatin accessibility profiles for *SHOX2*, *HMX2*, and *SIX3* motifs in control and mutant hypothalamus.

**(C)** Heatmap displaying key regulons affected in *Isl1* mutants across various hypothalamic regions.

**Figure S9. Disruption of *Nkx2.2* leads to altered hypothalamic regionalization and neuronal fate specification (related to Figure 5).**

**(A)** UMAP projections comparing hypothalamic progenitor populations in control and *Nkx2-2* mutant hypothalamus. Brighter colors represent higher cell density.

**(B)** Feature plots showing expression changes in key cell type-specific markers within *Nkx2-2* mutant.

**Figure S10. Dysregulation of transcription factors and motifs in *Dlx1/2* mutants (related to Figure 6).**

**(A)** Violin plots showing reduced expression of *Dlx1* and *Dlx2* expression in *Dlx1/2* mutants compared to controls at E12.5.

**(B)** Top: UMAP projections of hypothalamic progenitor populations in control and *Dlx1/2*-mutant hypothalamus. Bottom: Motif analysis showing altered accessibility of *DLX5*, *ISL1*, and *NR4A2* motifs in control versus mutant hypothalamus.

**(C)** Heatmap displaying key regulons in *Dlx1/2*-mutant hypothalamus compared to controls.

**Figure S11. Loss of *Dlx1/2* leads to downregulation of prethalamic markers and expansion of supramammillary nucleus (SMN)/medial mammillary nucleus (MMN) markers (related to Figure 6).**

**(A)** Feature plots from scRNA-Seq showing downregulation of prethalamic transcription factors *Arx*, *Isl1*, *Foxb1*, *Foxa1*, and *Hmx2* in *Dlx1/2* mutants compared to controls.

**(B)** *In situ* hybridization at E12.5 and E13.5 confirms changes observed in the scRNA-Seq dataset. Scale bar: 400 μm.

**(C)** Fluorescent *in situ* hybridization and immunostaining at E13.5 showing disrupted expression of prethalamic markers *Sp8*, *Meis2*, and Pax6 in *Dlx1/2*-mutants. *Pitx2*, a marker of the SMN/MMN, shows dorsal expansion into the prethalamic region in mutants, indicating a loss of repression of caudal fates. Scale bar: 400 μm.

**Figure S12. Loss of *Dlx1/2* disrupts hypothalamic neuronal identity, feeding behavior, and survival (related to Figure 7).**

**(A)** UMAP projections showing hypothalamic neuronal clusters at P8 in control and *Dlx1/2*-mutant mice.

**(B)** Feature plots (top row) and fluorescent *in situ* hybridization (bottom row) showing the loss of *Npw* expression in the DMH of *Dlx1/2*-mutant mice compared to controls at P8. Scale bar: 30 μm.

**(C)** Quantification of body weight and body length at P21 in control and *Dlx1/2*-mutant mice. Mutant mice show significant reductions in both weight (*\*\*\*\* p < 0.0001*) and length (*\*\*\*p < 0.001*).

**(D)** Kaplan-Meier survival curves showing the probability of survival in *Dlx1/2* mutants and controls on standard chow diet (top panel) and gel-based diet (bottom panel). Mutants show significantly reduced survival on both diets, with more rapid mortality on a chow diet (*p < 0.0001* for the chow diet, *p < 0.0005* for a gel-based diet).

**(E)** Feature plots showing loss of *Gal* in the Dlx1/2-positive regions but upregulation of *Gal* in the Dlx1/2-negative regions (thalamus) (top panel), and ectopic expression of *Arxes1* in the Dlx1/2-positive regions (bottom panel).

**(F)** Fluorescent *in situ* hybridization showing reduction of *Pnoc* in the TRN (top row), and ectopic expression of *Gal* in *Slc17a6-*expressing neurons in the thalamus (bottom row). Scale bar: 30 μm.

**Figure S13. Loss of *Dlx1/2* disrupts neuronal populations and increases gliosis in the TRN and ZI (related to Figure 7).**

**(A)** Top: schematic showing that conditional knockout of *Dlx1* and *Dlx2* in the developing prethalamus and hypothalamus was generated using the *Foxd1-Cre* driver to study the TRN and ZI. Bottom: predicted gene regulatory networks (GRNs) show interactions between *Dlx1* and *Dlx2*.

**(B)** snRNA-Seq analysis of TRN and ZI cell types. UMAP plots of control (left) and *Dlx1/2* mutant (right), highlighting altered neuronal clusters.

**(C)** Reduced parvalbumin (Pvalb)-expressing neurons in the TRN and ZI of *Dlx1/2* mutants: representative immunostaining for Pvalb (red) and DAPI (blue) in control (left) and *Dlx1/2* mutant (middle). Scale bar: 250 μm. Quantification (Right) shows a significant reduction in the percentage of Pvalb-expressing neurons in the mutant compared to the control. *** *p < 0.001*, unpaired t-tests.

**(D)** Spatial transcriptomics for *Gal* reveals ectopic expression in the thalamus.

**(E)** Reduced neuronal density in the TRN of Dlx1/2 mutants: Quantification of HuC/D-& NeuN-cells per mm² in the TRN. ** *p < 0.01*, unpaired t-tests.

**(F)** Gliosis in Dlx1/2 mutants: Spatial transcriptomics for *Opalin* (myelin marker, purple), *Gfap* (astrocytic marker, red), and *Cd68* (microglial activation marker, yellow) reveal increased gliosis in the mutant TRN (right) compared to Control (left). Dashed lines outline TRN boundaries. Scale bar: 250 μm.

**(G)** Miniature excitatory postsynaptic current (mEPSC) recordings from the medial geniculate nucleus (MGN), ventral posteromedial nucleus (VPM), and lateral geniculate nucleus (LGN) of control and *Dlx1/2* mutant mice. Left: Schematic of recording site and representative mEPSC traces from control and mutant. Right: Quantification of mEPSC parameters: Average frequency, Average amplitude, and Average decay time across 3 brain regions (top row) or pooled analysis (bottom row) from control (blue) and mutant (red). Data are presented as mean ± SEM, with individual data points shown.

**Figure S14. Behavioral deficits in *Dlx1/2* mutants (related to Figure 7).**

**(A)** Increased activity in *Dlx1/2* mutants with DeepLabCut: Left and middle panels show representative DeepLabCut movement heatmaps of Control (left) and Dlx1/2 mutant (middle) mice over a 1-hour trial. Color gradients indicate body position (nose to tail) over time. Right panels show quantification of total distance traveled (top panel) and time spent moving (bottom panel), both significantly increased in mutants compared to Control. \**p < 0.05*, unpaired t-tests.

**(B)** Exploratory behavior and locomotor patterns in the open field test: Quantification of the center-to-periphery ratio (top left panel) indicates decreased thigmotaxis in mutants compared to Control. Additional metrics show a significant increase in center distance traveled, time spent in the center, and the number of center entries for mutants. However, center’s average speed was not significantly altered. For the periphery, mutants display decreased periphery time, increased periphery entries, and increased periphery average speed. * *p < 0.05*, ** *p < 0.01*, unpaired t-tests.

**(C)** Sensory responses assessed by hot and cold plate tests: mutants exhibit a significant decrease in cold plate latency, indicative of altered sensory response to cold stimuli (* *p < 0.05*, unpaired t-tests). No significant differences are observed in the hot plate latency between control and mutants.

**(D)** Motor coordination assessed by the rotor rod test: Performance on the accelerating rotor rod over three consecutive days is shown for latency to fall (left) and rotational speed (revolutions per minute, RPM, at fall; right). Mutants exhibit no significant deficits compared to control in either measure, suggesting intact motor coordination and endurance.

## Tables

Table S1: Sample Information

Table S2: scRNA-Seq Data for Fig 1B

Table S3: scATAC-Seq Data for Fig 1B

Table S4: scRNA-Seq Data for Fig 1C

Table S5: scATAC-Seq Data for Fig 1C

Table S6: scRNA-Seq Data for Fig S1B

Table S7: scRNA-Seq Data for Fig S2A

Table S8: SCENIC+ Output for E11-E14

Table S9: Extended Data for Fig 2D

Table S10: SCENIC+ Output for E15-P8

Table S11: Extended Data for Fig S6

Table S12: scRNA-Seq Data for Fig 4B, D

Table S13: Extended Data for Fig 4C

Table S14: scRNA-Seq Data for Fig 4F and 4H

Table S15: SCENIC+ Output for Isl1 Mutants

Table S16: Extended Data for Fig 4G

Table S17: scRNA-Seq Data for Fig 5B

Table S18: scRNA-Seq Data for Fig 5C and 5D

Table S19: SCENIC+ Output for E12.5 Dlx1/2 Mutants

Table S20: scRNA-Seq Data for Fig 6

Table S21: Extended Data for Fig 6C

Table S22: Extended Data for Fig S12B

Table S23: Extended Data for Fig S12C

Table S24: Extended Data for Fig S12D, Standard chow diet

Table S25: Extended Data for Fig S12D, Gel-based food

Table S26: Extended Data for Fig S13B, Proportion of snRNA-Seq clusters

Table S27: Extended Data for Fig S13B, differential genes across snRNA-Seq clusters

Table S28: Xenium Probe Specifications

Table S29: Xenium Observations

Table S30: DeepLabCut Output Control representative data

Table S31: DeepLabCut Output Dlx1/2 Mutant representative data

Table S32: Open Field Test Raw Data

Table S33: Hot Plate Test Raw Data

Table S34: Cold Plate Test Raw Data

Table S35: Rotarod Test Raw Data

Table S36: Slice electrophysiology - Group Information

Table S37: Slice electrophysiology - Excitation Frequency

Table S38: Slice electrophysiology - Inhibition Frequency

Table S39: Slice electrophysiology - Excitation Amplitude

Table S40: Slice electrophysiology - Inhibition Amplitude

Table S41: Slice electrophysiology - Excitation Decay Time

Table S42: Slice electrophysiology - Inhibition Decay Time

Table S43: Slice electrophysiology - Mean and SEM

## Abbreviations

ARC: Arcuate Nucleus
CNS: Central Nervous System
DMH: Dorsomedial Hypothalamus
E: Embryonic Day
EmThal: Thalamic Eminence
GRNs: Gene Regulatory Networks
GWAS: Genome-Wide Association Study
ID: Intrahypothalamic Diagonal
LGN: Lateral Geniculate Nucleus
LH: Lateral Hypothalamus
mEPSCs: Miniature Excitatory Postsynaptic Currents
MGN: Medial Geniculate Nucleus
mIPSCs: Miniature Inhibitory Postsynaptic Currents
MMN: Medial Mammillary Nucleus
NPC: Neural Progenitor Cell
NSC: Neural Stem Cell
P: Postnatal Day
PMN: Premammillary Nucleus
PreThal: Prethalamus
PVH: Paraventricular Nucleus of the Hypothalamus
SCN: Suprachiasmatic Nucleus
scATAC-Seq: Single-cell ATAC Sequencing
scRNA-Seq: Single-cell RNA Sequencing
SMN: Supramammillary Nucleus
SON: Supraoptic Nucleus
TFs: Transcription Factors
TRN: Thalamic Reticular Nucleus
Tub: Tuberal Hypothalamus
VMH: Ventromedial Hypothalamus
VPM: Ventral Posteromedial Nucleus
ZLI: Zona Limitans Intrathalamica
ZI: Zona Incerta

