## Supplemental Figures S1-14 for "Decoding Gene Networks Controlling Hypothalamic and Prethalamic Neuron Development"

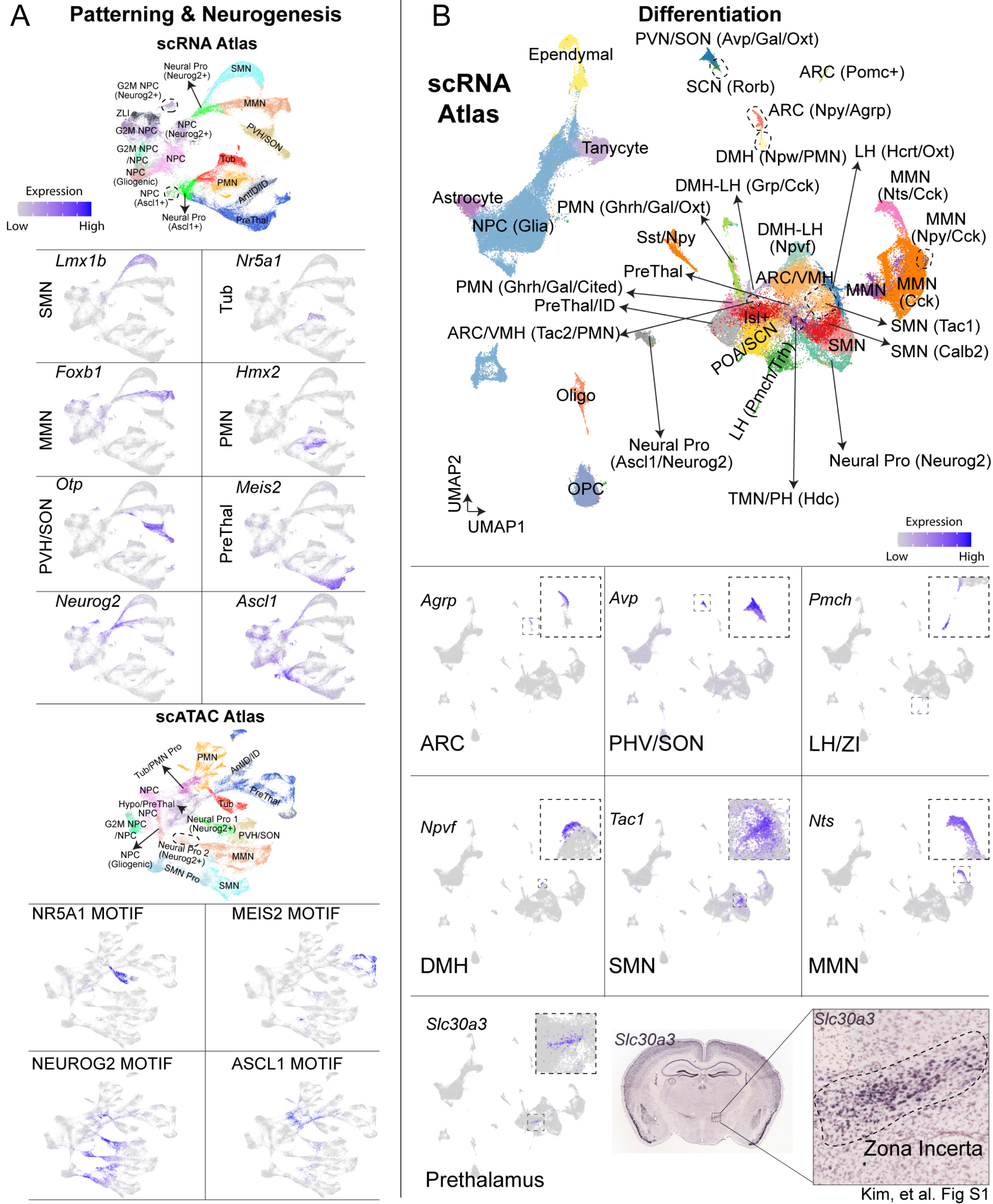
A
Patterning & Neurogenesis
scRNA Atlas
Neural Pro (Neurog2+)
SMN
G2M NPC (Neurog2+)
MMN
ZLI
NPC (Neurog2+)
G2M NPC
PVH/SON
G2M NPC /NPC
NPC
Tub
NPC (Gliogenic)
PMN
NPC (Ascl1+)
AntID/ID
Neural Pro (Ascl1+)
PreThal
Expression
Low
High
Lmx1b
SMN
Nr5a1
Tub
Foxb1
MMN
Hmx2
PMN
Otp
PVH/SON
Meis2
PreThal
Neurog2
Ascl1
scATAC Atlas
Tub/PMN Pro
PMN
AntID/ID
NPC
PreThal
Hypo/PreThal NPC
Tub
G2M NPC /NPC
Neural Pro 1 (Neurog2+)
PVH/SON
NPC (Gliogenic)
Neural Pro 2 (Neurog2+)
MMN
SMN Pro
SMN
NR5A1 MOTIF
MEIS2 MOTIF
NEUROG2 MOTIF
ASCL1 MOTIF
B
Differentiation
scRNA Atlas
Ependymal
PVN/SON (Avp/Gal/Oxt)
SCN (Rorb)
ARC (Pomc+)
ARC (Npy/Agrp)
Tanycyte
DMH (Npw/PMN)
LH (Hcrt/Oxt)
DMH-LH (Grp/Cck)
MMN (Nts/Cck)
Astrocyte
PMN (Ghrh/Gal/Oxt)
MMN (Npy/Cck)
NPC (Glia)
Sst/Npy
DMH-LH (Npvf)
PreThal
ARC/VMH
MMN
MMN (Cck)
PMN (Ghrh/Gal/Cited)
PreThal/ID
Isl+
SMN (Tac1)
ARC/VMH (Tac2/PMN)
POA/SCN
SMN
SMN (Calb2)
LH (Pmch/Trh)
Oligo
Neural Pro (Ascl1/Neurog2)
Neural Pro (Neurog2)
UMAP2
UMAP1
OPC
TMN/PH (Hdc)
Expression
Low
High
Agrp
ARC
Avp
PHV/SON
Pmch
LH/ZI
Npvf
DMH
Tac1
SMN
Nts
MMN
Slc30a3
Prethalamus
Slc30a3
Slc30a3
Zona Incerta
Kim, et al. Fig S1

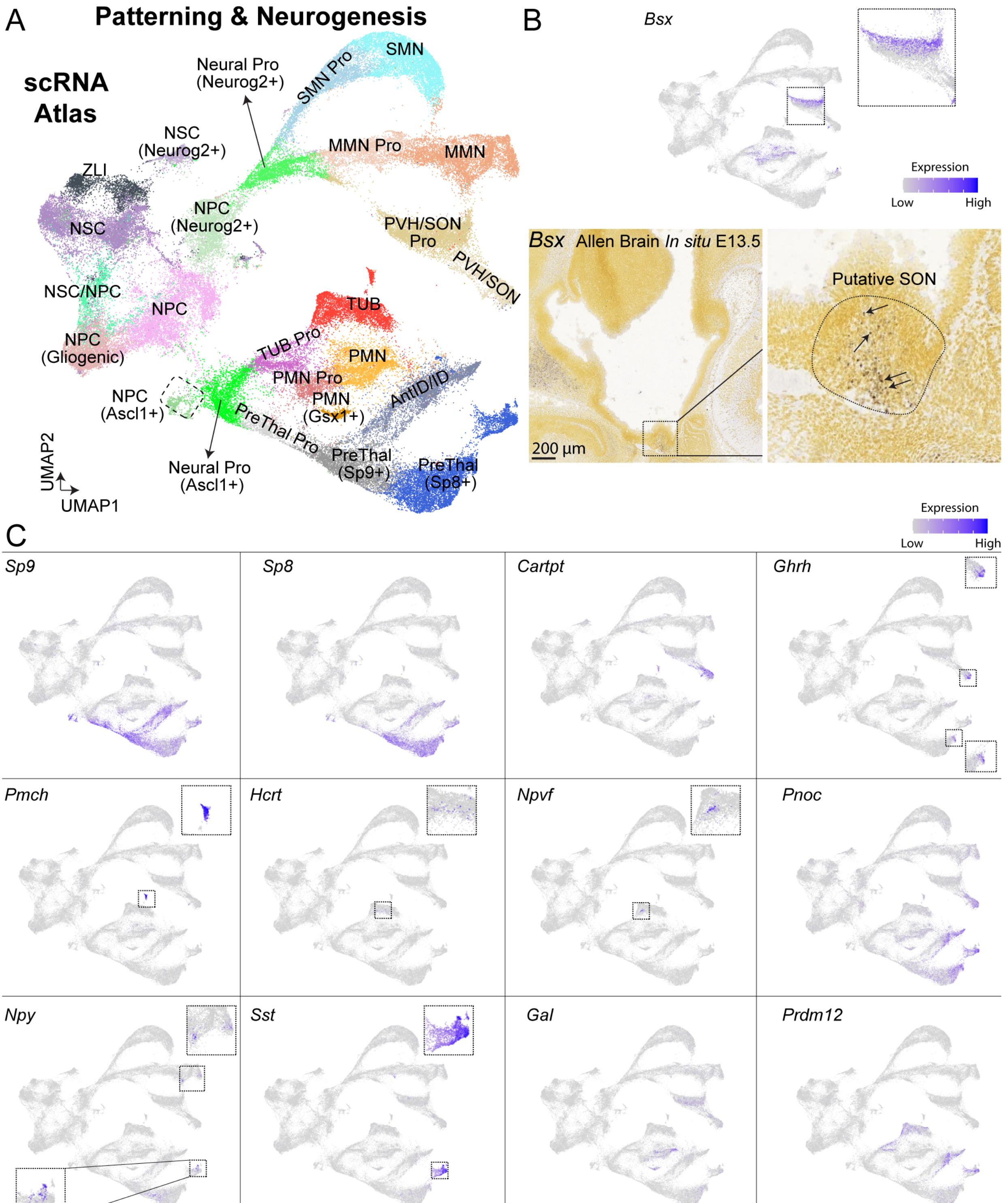


Kim, et al. Fig S2

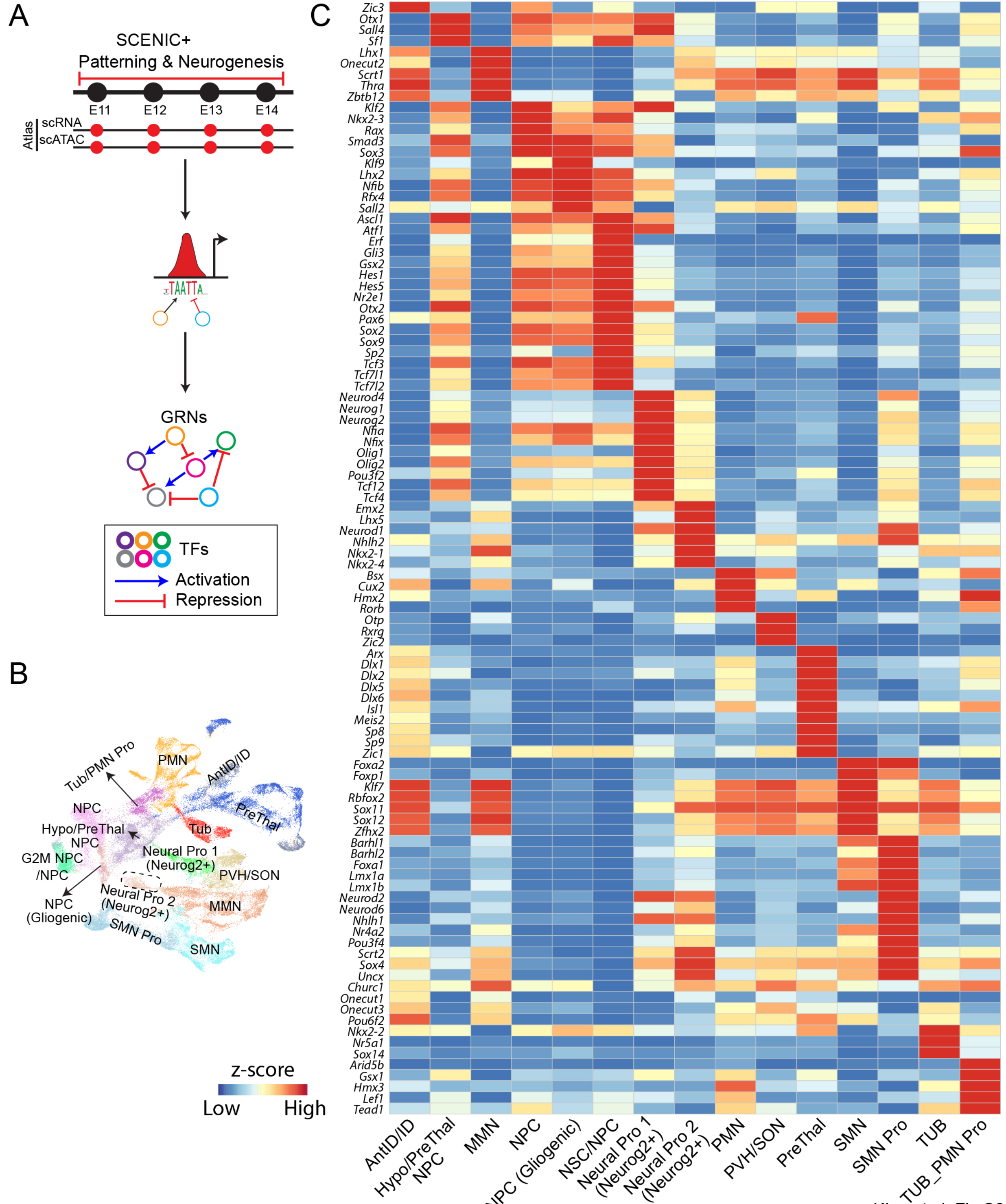


Kim, et al. Fig S3

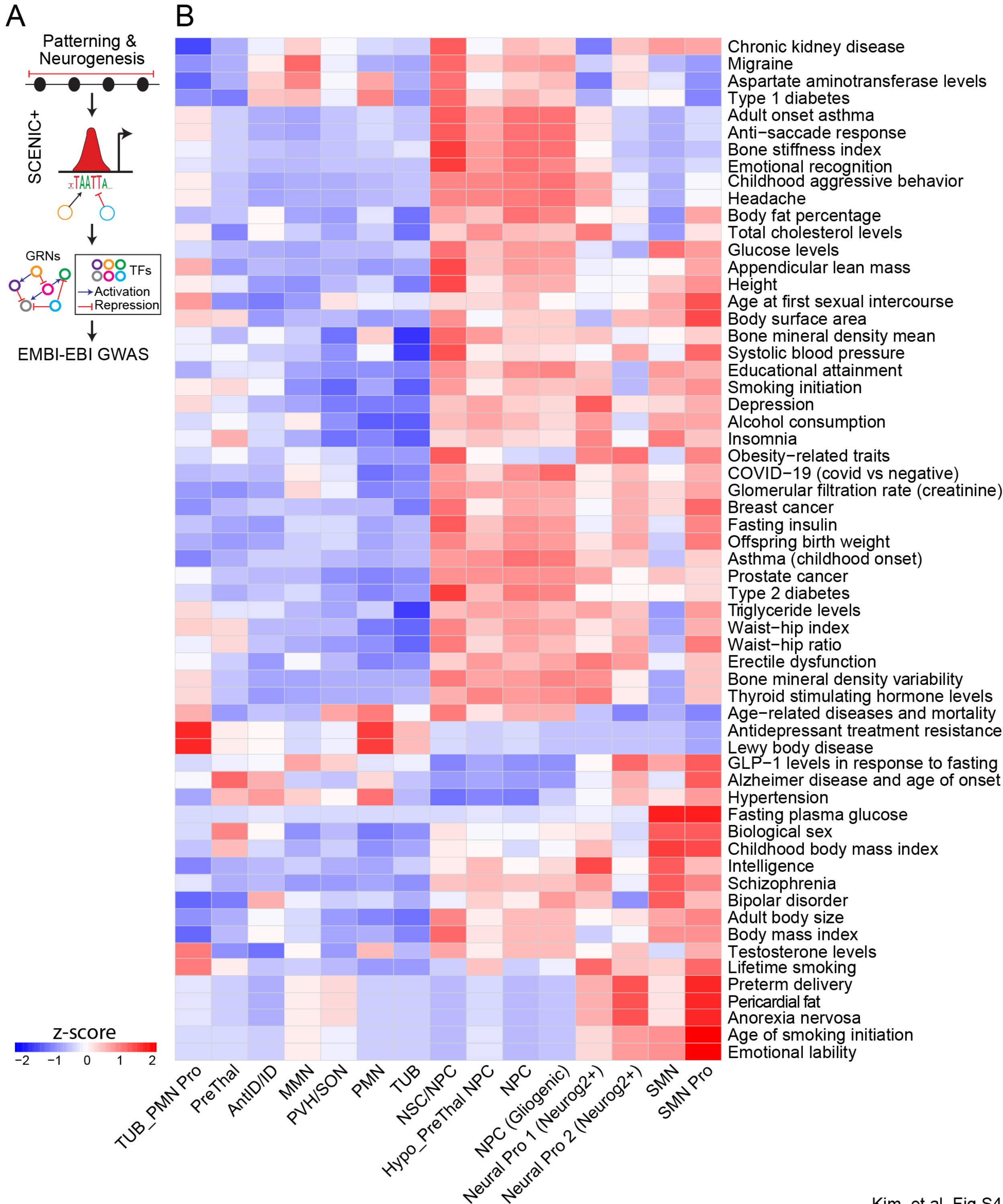


Kim, et al. Fig S4

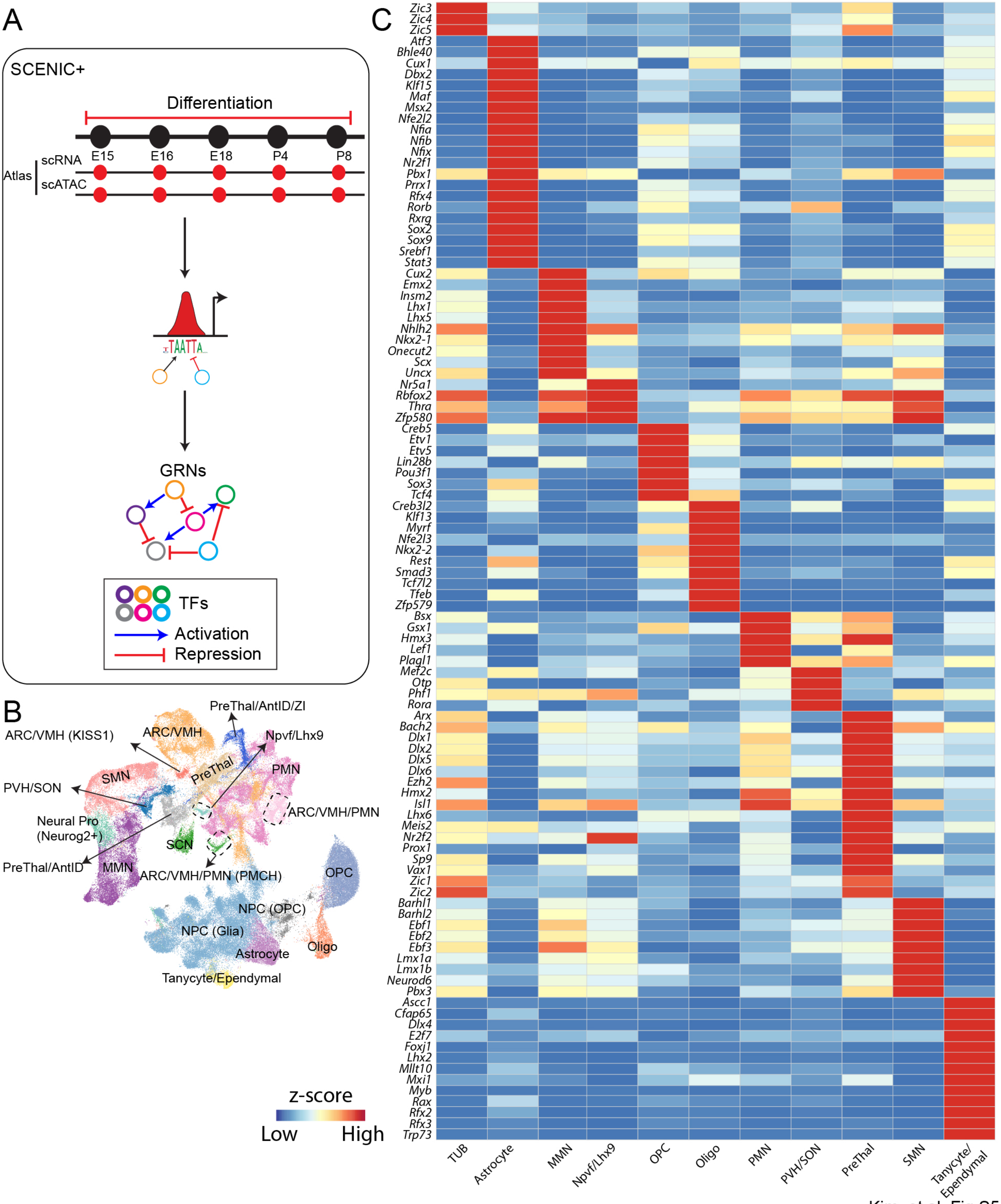


Kim, et al. Fig S5

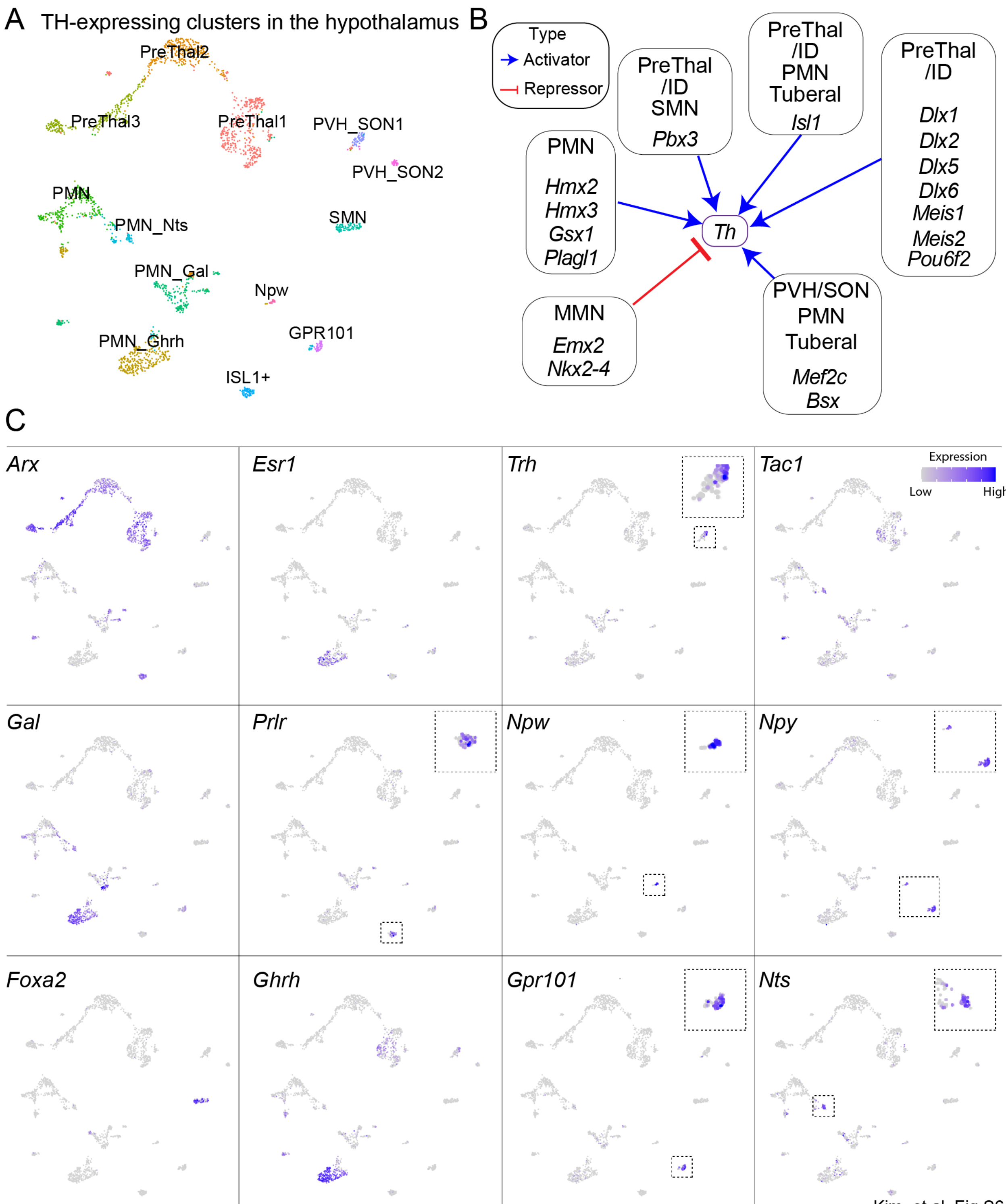


Kim, et al. Fig S6

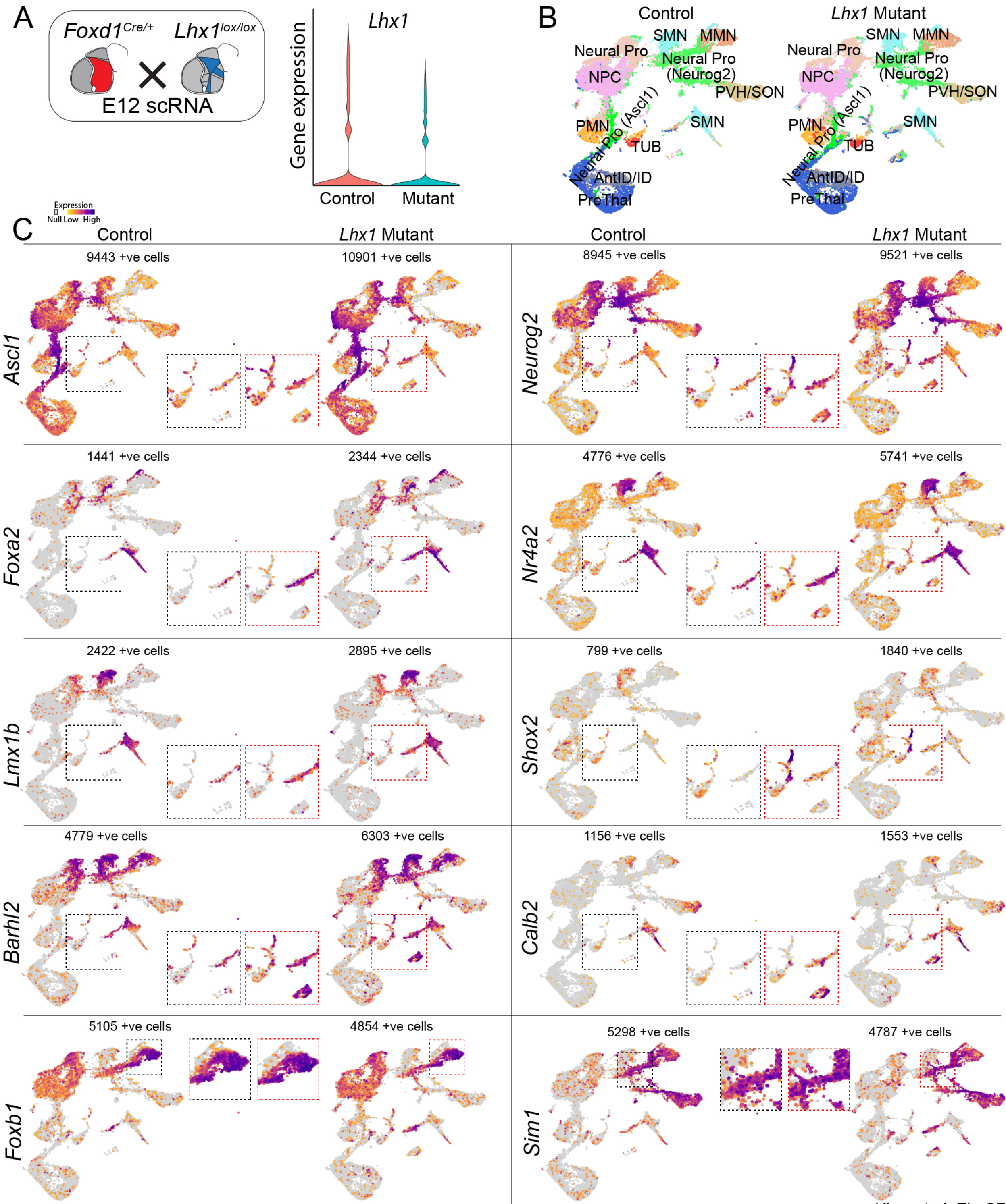


Kim, et al. Fig S7

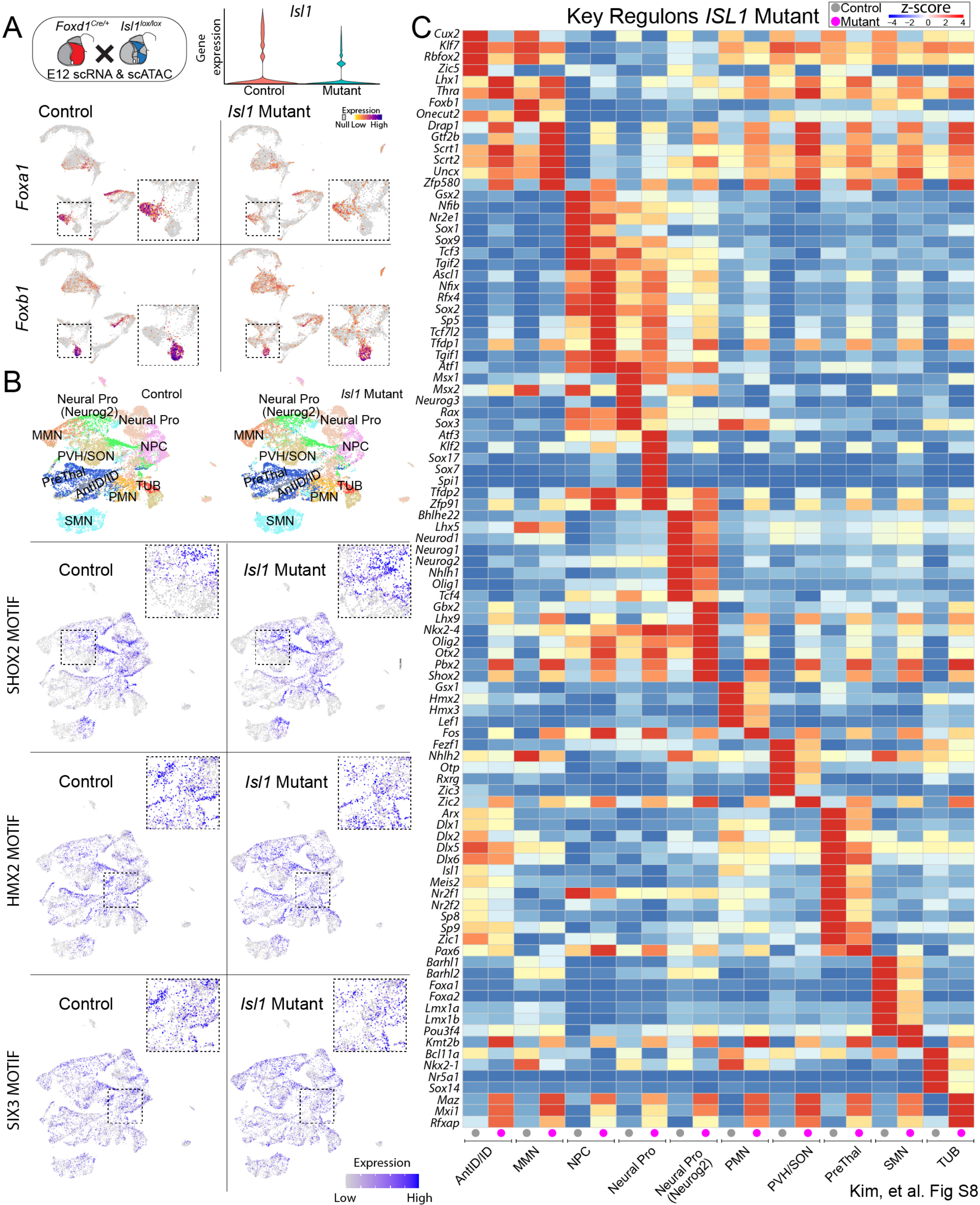


Kim, et al. Fig S8

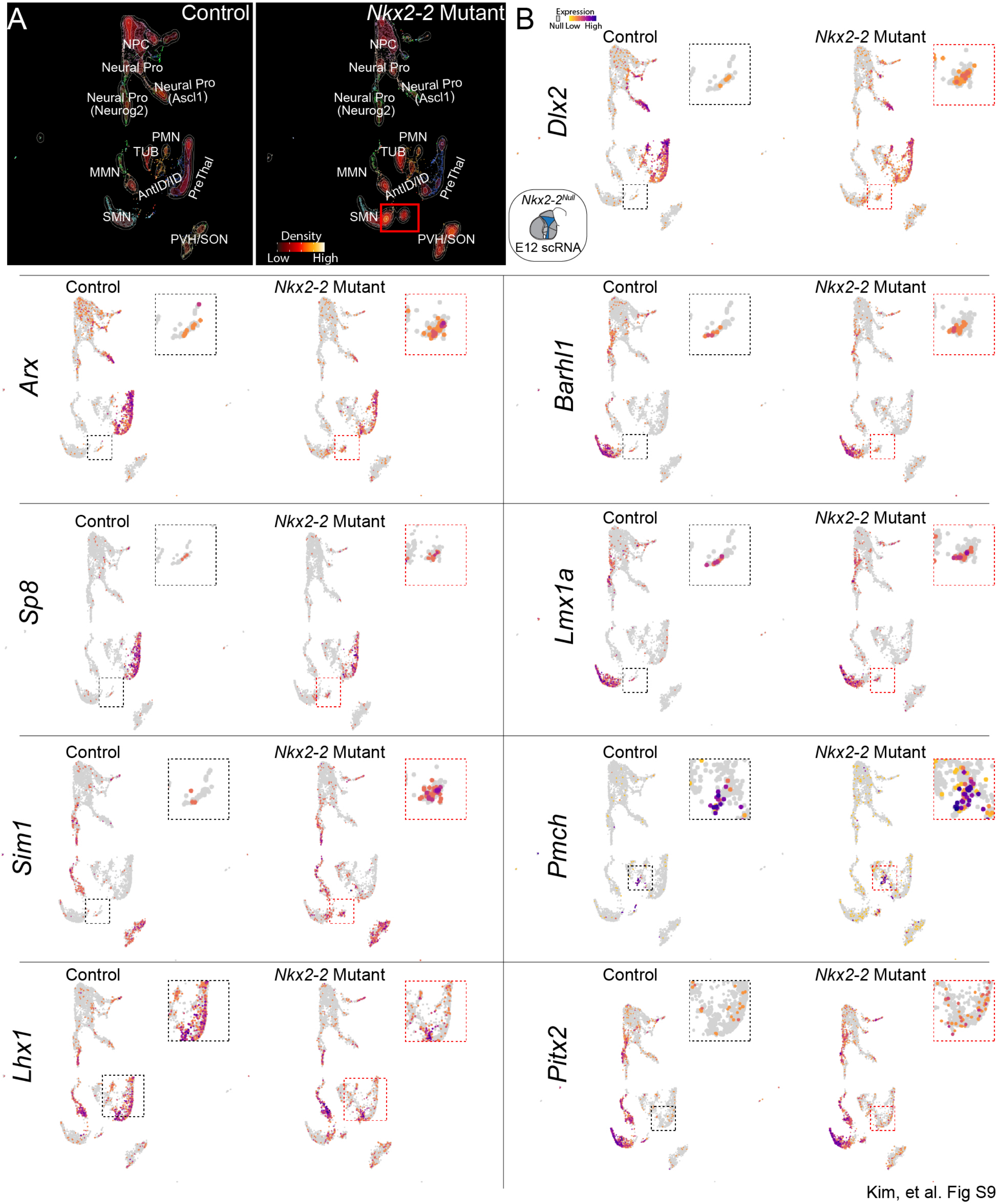


Kim, et al. Fig S9

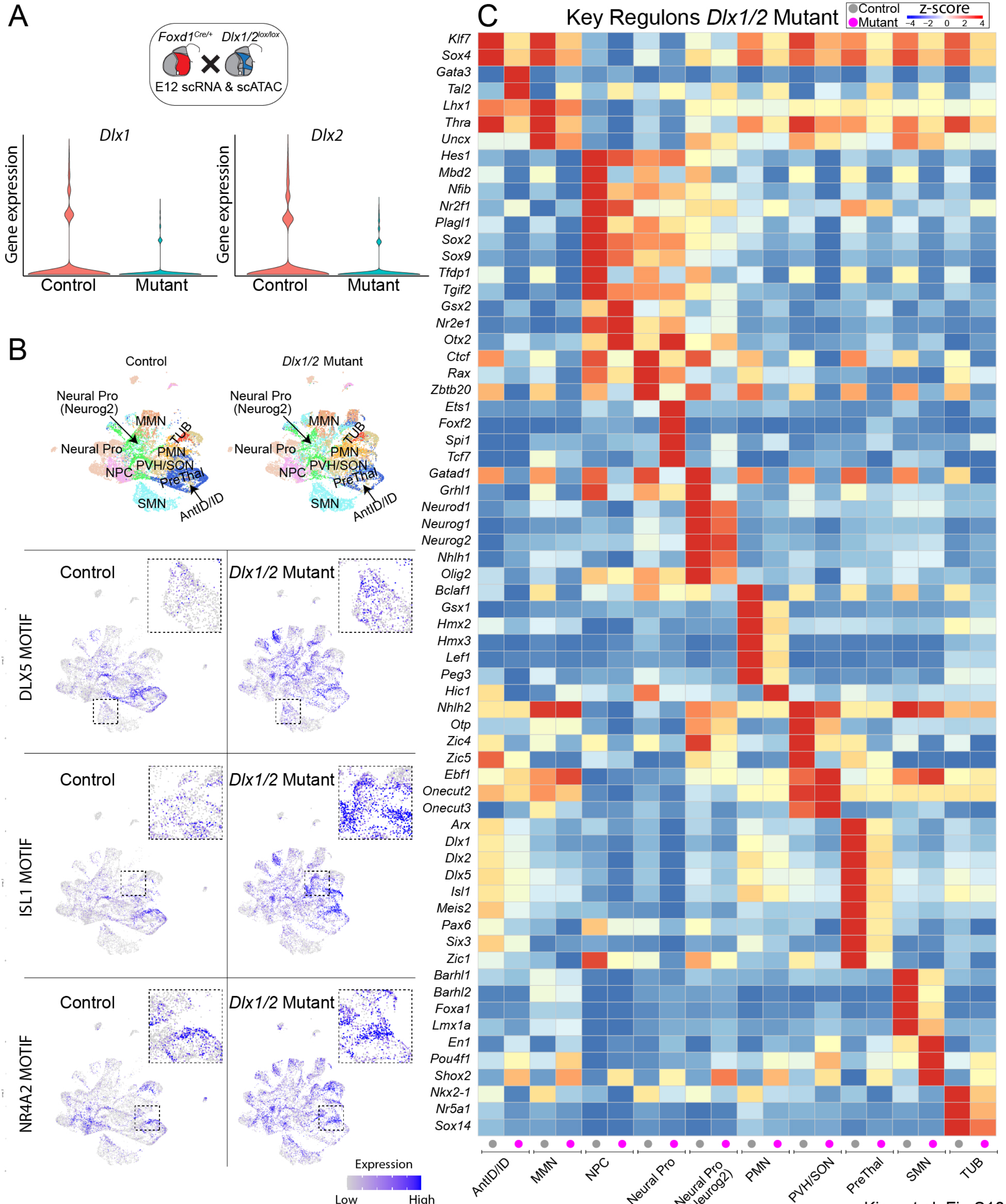


Kim, et al. Fig S10

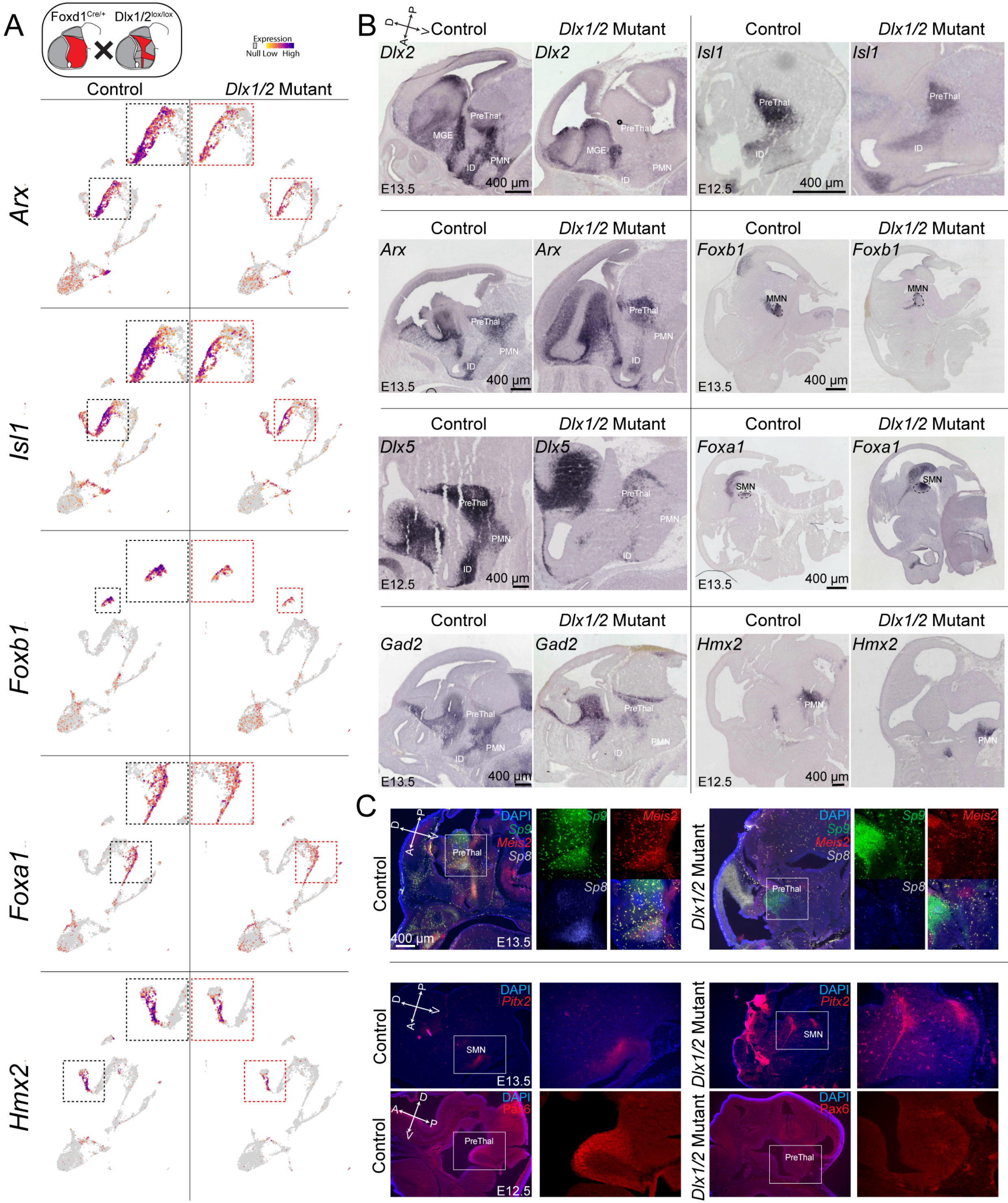


Kim, et al. Fig S11

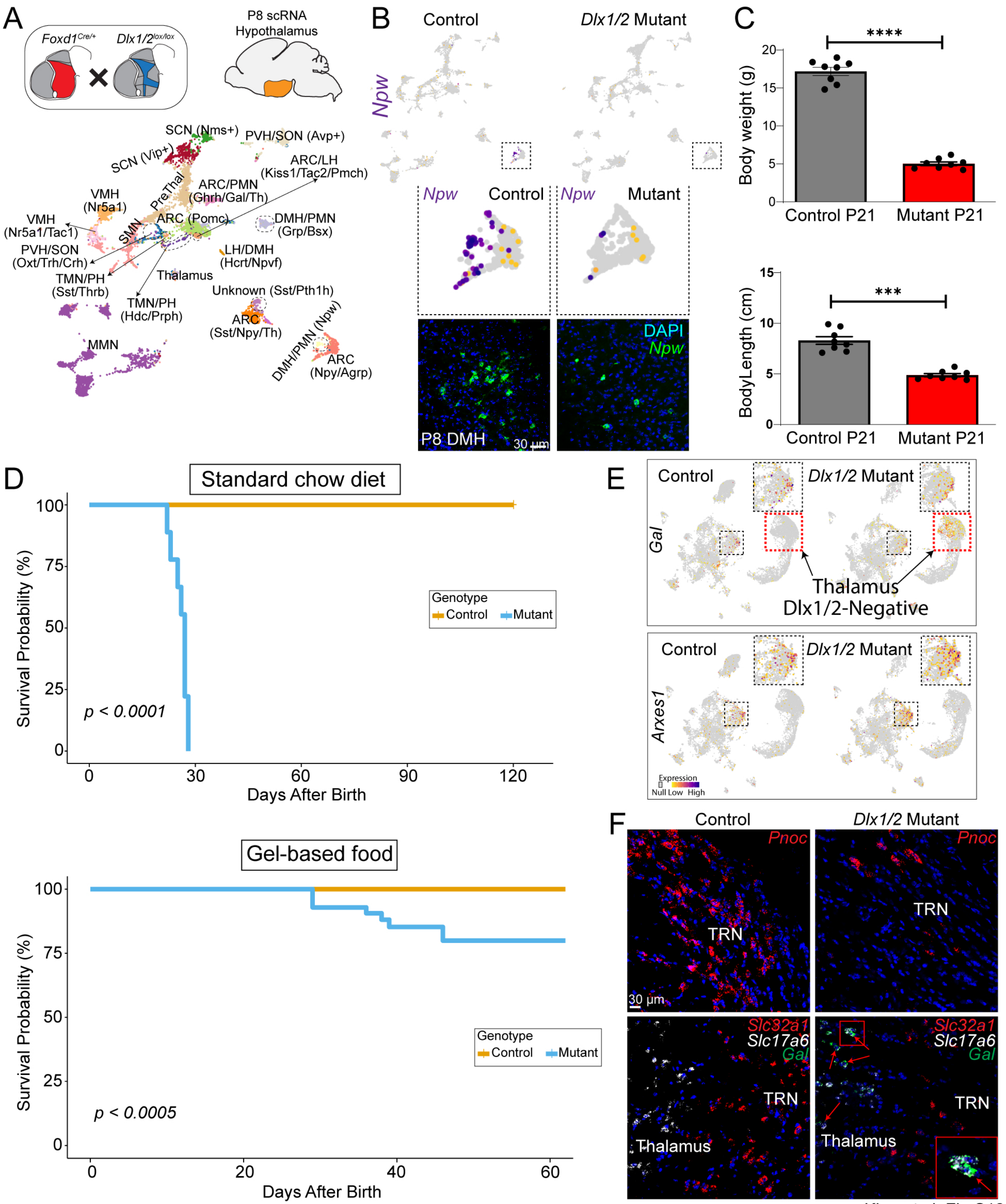
A
Foxd1Cre/+
Dlx1/2lox/lox
P8 scRNA
Hypothalamus
SCN (Nms+)
PVH/SON (Avp+)
SCN (Vip+)
ARC/LH
(Kiss1/Tac2/Pmch)
PreThal
ARC/PMN
(Ghrh/Gal/Th)
VMH
(Nr5a1)
VMH
(Nr5a1/Tac1)
SMN
ARC (Pomc)
DMH/PMN
(Grp/Bsx)
PVH/SON
(Oxt/Trh/Crh)
LH/DMH
(Hcrt/Npvf)
TMN/PH
(Sst/Thrb)
Thalamus
TMN/PH
(Hdc/Prph)
Unknown (Sst/Pth1h)
ARC
(Sst/Npy/Th)
DMH/PMN (Npw)
ARC
(Npy/Agrp)
MMN
B
Control
Dlx1/2 Mutant
Npw
Npw
Control
Npw
Mutant
DAPI
Npw
P8 DMH
30 μm
C
****
Body weight (g)
Control P21
Mutant P21
***
BodyLength (cm)
Control P21
Mutant P21
D
Standard chow diet
Survival Probability (%)
Genotype
Control
Mutant
p < 0.0001
Days After Birth
Gel-based food
Survival Probability (%)
Genotype
Control
Mutant
p < 0.0005
Days After Birth
E
Control
Dlx1/2 Mutant
Gal
Thalamus
Dlx1/2-Negative
Control
Dlx1/2 Mutant
Arxes1
Expression
Null Low High
F
Control
Dlx1/2 Mutant
Pnoc
TRN
Pnoc
TRN
30 μm
Slc32a1
Slc17a6
Gal
TRN
Thalamus
Slc32a1
Slc17a6
Gal
TRN
Thalamus

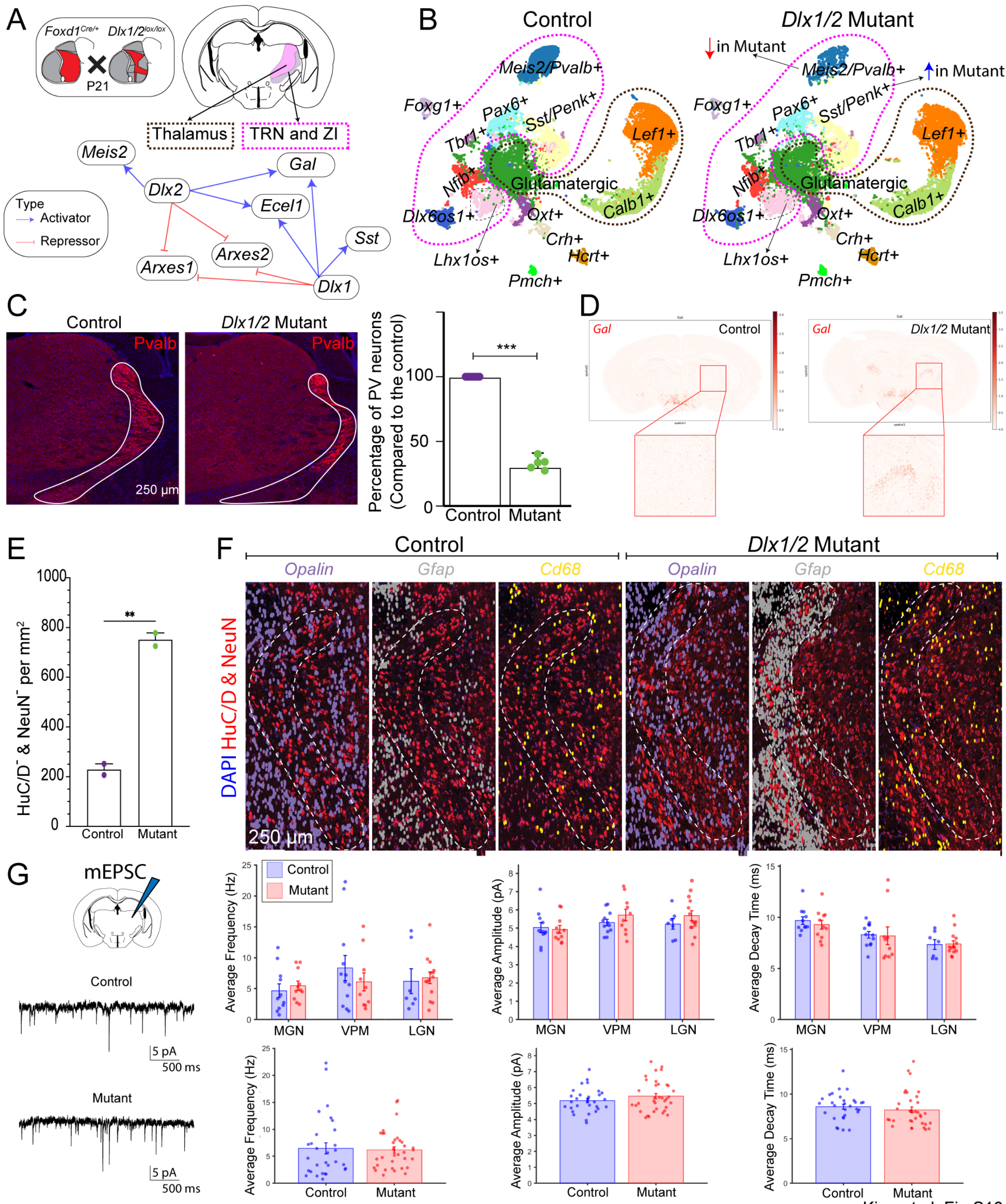


Kim, et al. Fig S13

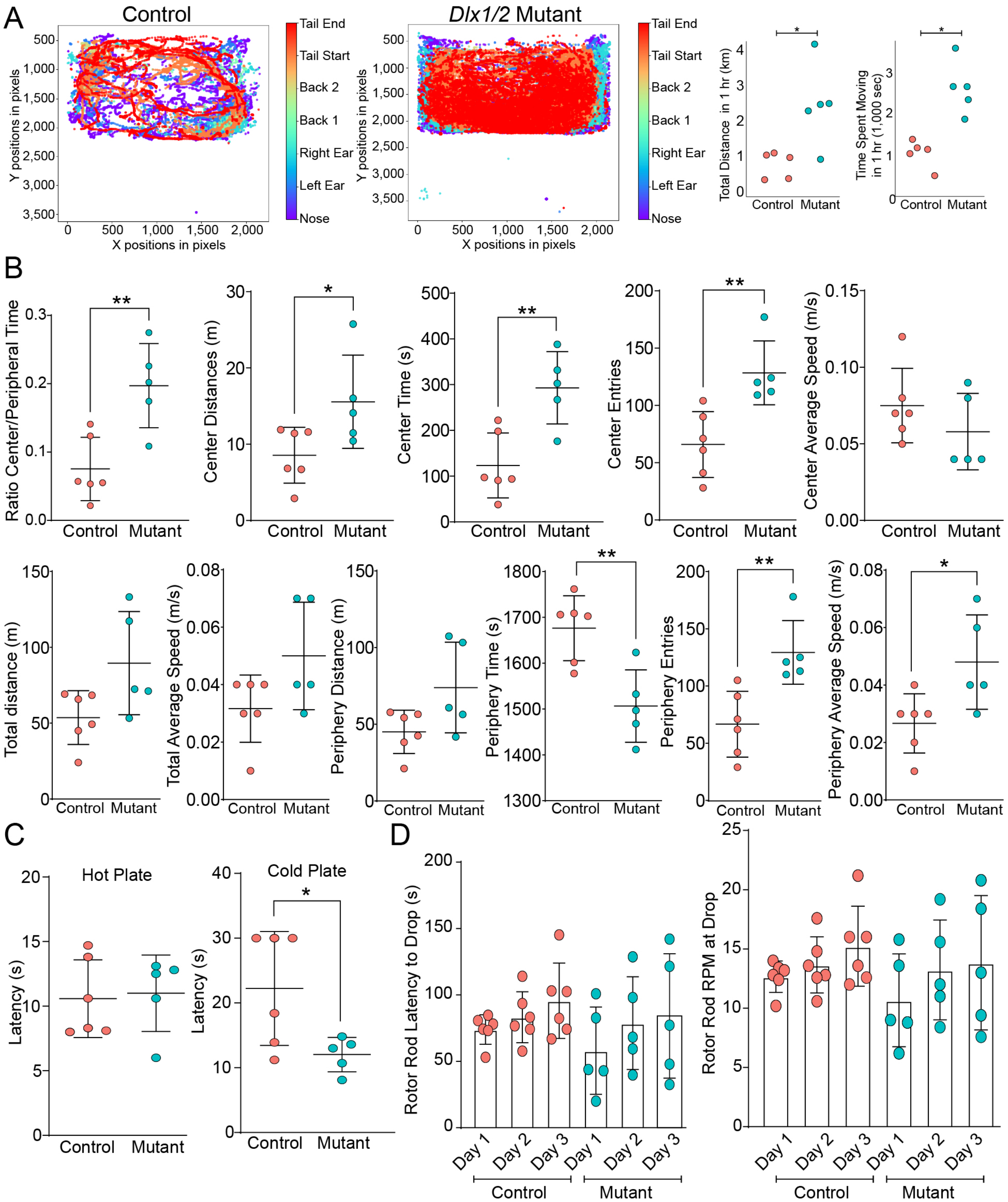

Kim, et al. Fig S14
